# The Arabidopsis dehydrin ERD14 chaperones the brassinosteroid receptor kinase BRL3 to the plasma membrane to confer drought resistance

**DOI:** 10.1101/2025.01.15.633155

**Authors:** M.M. Marquès-Bueno, I. Herrero-García, S. Pistillo, A. Gupta, F. Lozano-Elena, D Blasco-Escámez, M. Karampelias, N. Fàbregas, V.I. Strotmann, Y. Stahl, A.I Caño-Delgado

## Abstract

Brassinosteroid plant hormones are involved in growth, biotic and abiotic-stress signaling, but functional analysis of its multiple receptors has largely focused on the ubiquitously expressed BRASSINOSTEROID INSENSITIVE 1 (BR1). The vascular-expressed brassinosteroid receptor kinase BRI1-LIKE 3 (BRL3) has a specific and essential role in Arabidopsis adaptation to drought and elevated temperature. However, the mechanistic understanding of BRL3 function and regulation is scant. Here, we show the dehydrin ERD14 colocalizes with BRI1 and BRL3 at the plasma membrane and physically interacts with these receptors through the ERD14^Lys^ domain for proper receptor recruitment. *erd14* mutants internalize BRL3–GFP and revert drought-resistant *BRL3* over-expression phenotypes. BRL3 activation and downstream signaling events, including dephosphorylation of BES1 transcription factor, depend on the chaperone function of ERD14. Our study shows how proper chaperoning by ERD14 regulates BRL3 subcellular localization and activity of the vascular receptor, prompting re-examination of models of brassinosteroid-receptor functional diversification in plant adaptation to abiotic stress from specific cells of the vascular systems.

## Introduction

Brassinosteroid hormones are essential regulators of plant growth and development, and necessary for plant adaptation to abiotic stress (Yao et al. 2023). Functional studies in Arabidopsis have shown that brassinosteroids are sensed by a 70-amino-acid island domain at the extracellular domain of the BRASSINOSTEROID INSENSITIVE 1 (BRI1) family of leucine-rich repeat receptor-like kinases (LRR–RLKs) (Kinoshita et al. 2005; Sun et al. 2010; Hothorn et al. 2011). The BRI1 receptor promotes growth and development from the outer cell layers (Savaldi-Goldstein et al. 2007; Kelly-Bellow et al. 2023), with Arabidopsis *bri1* mutants dwarf and sterile (Li and Chory 1997). BRI1 localizes to the plasma membrane and endomembranes (Geldner et al. 2007). In the absence of brassinosteroid ligand, BRI1 physically interacts with BRI1 KINASE INHIBITOR 1 (BKI1), thereby preventing interactions with the co-receptor BRASSINOSTEROID-INSENSITIVE-1-ASSOCIATED RECEPTOR KINASE 1 (BAK1). Upon treatment with the ligand brassinolide (BL), the most physiologically active of the brassinosteroids, BRI1 phosphorylates BKI1, which is then released from the plasma membrane to the cytoplasm (Wang and Chory 2006; Jaillais et al. 2011). This release allows BRI1 to interact with BAK1, also known as SERK3 (SOMATIC-EMBRYOGENESIS RECEPTOR KINASE 3), which in turn triggers an intracellular phosphorylation cascade (Li and Nam 2002; Russinova et al. 2004). This signaling pathway culminates in the activation and stabilization of the transcription factors BES1 (BRI1-EMS-SUPPRESSOR 1) (Yin et al. 2002) and BZR1 (BRASSINAZOLE RESISTANT 1) (Wang and Chory 2006), which directly regulate brassinosteroid-responsive genes to fine-tune development across different tissues.

Beyond BRI1, two other brassinosteroid receptors BRI1-like 1 (BRL1) and BRL3, are also known to bind brassinolide ligand (Caño-Delgado et al. 2004; Kinoshita et al. 2005; Planas-Riverola et al. 2019). BRLs are receptors expressed in the stem-cell niche and vasculature of root and shoot tissues and are associated with adaptation to environmental stress (Fàbregas et al. 2018; Lozano-Elena and Caño-Delgado 2019; Planas-Riverola et al. 2019). Given that brassinosteroids are cell–cell mobile via plasmodesmata, it is plausible to hypothesize spatially separate roles for BRI1 and BRL3 in growth and abiotic-stress adaptation, respectively (Wang and Chory 2006; Sun et al. 2010; Planas-Riverola et al. 2019; Benitez-Alfonso and Caño-Delgado 2023). The biochemical characterization of BRL3 receptor complexes by immunoprecipitation (IP) and liquid chromatography–tandem mass spectrometry (LC–MS/MS) has identified candidate BRL3-interacting proteins (Fàbregas et al. 2013). Proteins related to brassinosteroid function such as BAK1 (Li and Nam 2002; Li et al. 2002), with protein trafficking functions such as VHA-A2 (Dettmer et al. 2006), and some cytoplasmic signaling components such as CDPK6 were identified, supporting the role of BRL3 receptors in brassinosteroid signaling, although Arabidopsis *brl3-2* mutants are not dwarf as most biosynthetic and *bri1* signal-transduction mutants.

Emerging evidence indicates specific roles for BRL3 receptor in adaptation to drought and elevated temperatures (Fàbregas et al. 2018; Gupta et al. 2023). This property is likely from over-accumulation of osmoprotectant metabolites in the root (Caño-Delgado et al. 2004; Planas-Riverola et al. 2019). Despite the assumption of the BRLs performing similar functions due to their high homology and conservation (Caño-Delgado et al. 2004), emerging evidence implicates non-overlapping roles of BRI1 and BRL3 in adapting to abiotic stress (Fàbregas et al. 2018; Lozano-Elena and Caño-Delgado 2019; Planas-Riverola et al. 2019; Gupta et al. 2023), evidenced by Arabidopsis plants over-expressing BRI1 being susceptible to drought (Fontanet-Manzaneque et al. 2024), in contrast with severe drought resistance provided by the BRL3 overexpression (Fàbregas et al. 2018). The brassinosteroid-deficient and insensitive mutant phenotypes associated with adaptation mechanisms such as hydrotropism or drought resistance may be related to severe developmental defects that lead to the dwarfism they exhibit (Takahashi et al. 2002; Yuan et al. 2018; Hola 2019; Xiang et al. 2021).

The stability of the brassinosteroid receptors in the membrane is related to downstream signaling (Irani et al. 2012). Upon ubiquitination, BRI1 is targeted for degradation, and it is detached from the plasma membrane, unable to induce signaling (Martins et al. 2015). Moreover, BRI1 is stabilized in the plasma membrane by the chaperone-like DE-ETIOLATION IN THE DARK AND YELLOWING IN THE LIGHT (DAY), regulating photomorphogenesis (Li et al. 2015; Lee et al. 2021).

Here, we sought to mechanistically characterize BRL3 function and regulation in Arabidopsis drought signaling through analysis of its interacting protein partners. Analysis of the BRL3 interactome (Fàbregas et al. 2018) revealed ERD14 as a promising candidate to control the stability and signaling of BRL3 upon abiotic stress for at least three main reasons. Firstly, ERD14 is highly induced upon abiotic stress (Kiyosue et al. 1994; Nylander et al. 2001). Secondly, it acts as chaperone to prevent heat-induced aggregation and/or inactivation of several substrates (Kovacs et al. 2008; Klimecka et al. 2020; Nguyen et al. 2020) and lastly, its expression is enriched in the vascular tissue (Winter et al. 2007). ERD14 belongs to the Dehydrin protein family, which is a group of non-catalytic proteins with protective functions involved in abiotic-stress resistance. They are defined by four highly conserved domains. Firstly, the K-segment (EKKGIMDKIKEKLPG), responsible for plasma-membrane localization (Allagulova et al. 2003; Hara 2010; Hanin et al. 2011); the S-segment, a serine-rich region that, upon phosphorylation, is responsible for nuclear localization (Hanin et al. 2011; Klimecka et al. 2020); the F-segment, a lysine-rich region related to cryoprotective properties (Richard Strimbeck 2017; Ohkubo et al. 2020), and lastly the charged-peptide segment (ChP-segment), also called the KEKE-segment, with homology with classical protein chaperones and nucleotide binding (Graether and Boddington 2014; Covarrubias et al. 2017; Murvai et al. 2021). Both K-segment and ChP-segment are lysine rich domains that conform a poly-lysine rich region in ERD14.

In this study we show BRL3 physically interacts with ERD14 through its poly-lysine domain to promote drought-stress tolerance. This interaction takes place at the plasma membrane, wherein ERD14 maintains and stabilizes BRL3 abundance by preventing receptor ubiquitination to enable proper spatial brassinosteroid signaling upon stress. Activation of BES1 is dependent on ERD14. *erd14* mutants suppress the phenotype of *BRL3* over-expression, highlighting a major role for the BRL3 co-receptor complex in regulation of abiotic-stress adaptation. Our study details the molecular basis of BRL3 function in adaptation and prompts re-examination of the functional diversification and physiological relevance of previously overlooked brassinosteroid receptors.

## Results

### ERD14 promotes BRL3 stability and activity on the plasma membrane by preventing ubiquitination

Over-expression of BRL3–GFP up-regulates brassinosteroid-biosynthesis genes and downstream BES1 and BZR1 effectors in Arabidopsis (Fàbregas et al. 2018). ERD14 was identified as a potential interactor in BRL3 complexes by mass spectrometry following co-immunoprecipitation (Fàbregas et al. 2013). Toward validating and understanding the functional and physiological relevance of the candidate BRL3–ERD14 complex in brassinosteroid ligand perception and downstream signaling, we measured root length of 7-d-old Arabidopsis seedlings following treatment with 0.4 nM brassinolide (BL) for 3 d that reduces root growth of plants able to perceive and signal the ligand (Figure 1A,B). *brl3-2, erd14-1* and *erd14-3* single mutants and the double *brl3-2 erd14-3* mutant were less sensitive than wild-type Col-0 controls. As reported for BRI1 (Wolf et al. 2014), over-expression of BRL3–GFP saw increased sensitivity to brassinolide, resulting in a stronger root-growth inhibition, as shown in the growth ratios between treatment and control (Figure 1B). This response for *35S_pro_:BRL3–GFP* plants was abolished to WT levels in the presence of *erd14-3*, thereby confirming the involvement of ERD14 in the brassinosteroid signaling pathway (Figure 1B).

**Figure 1.**
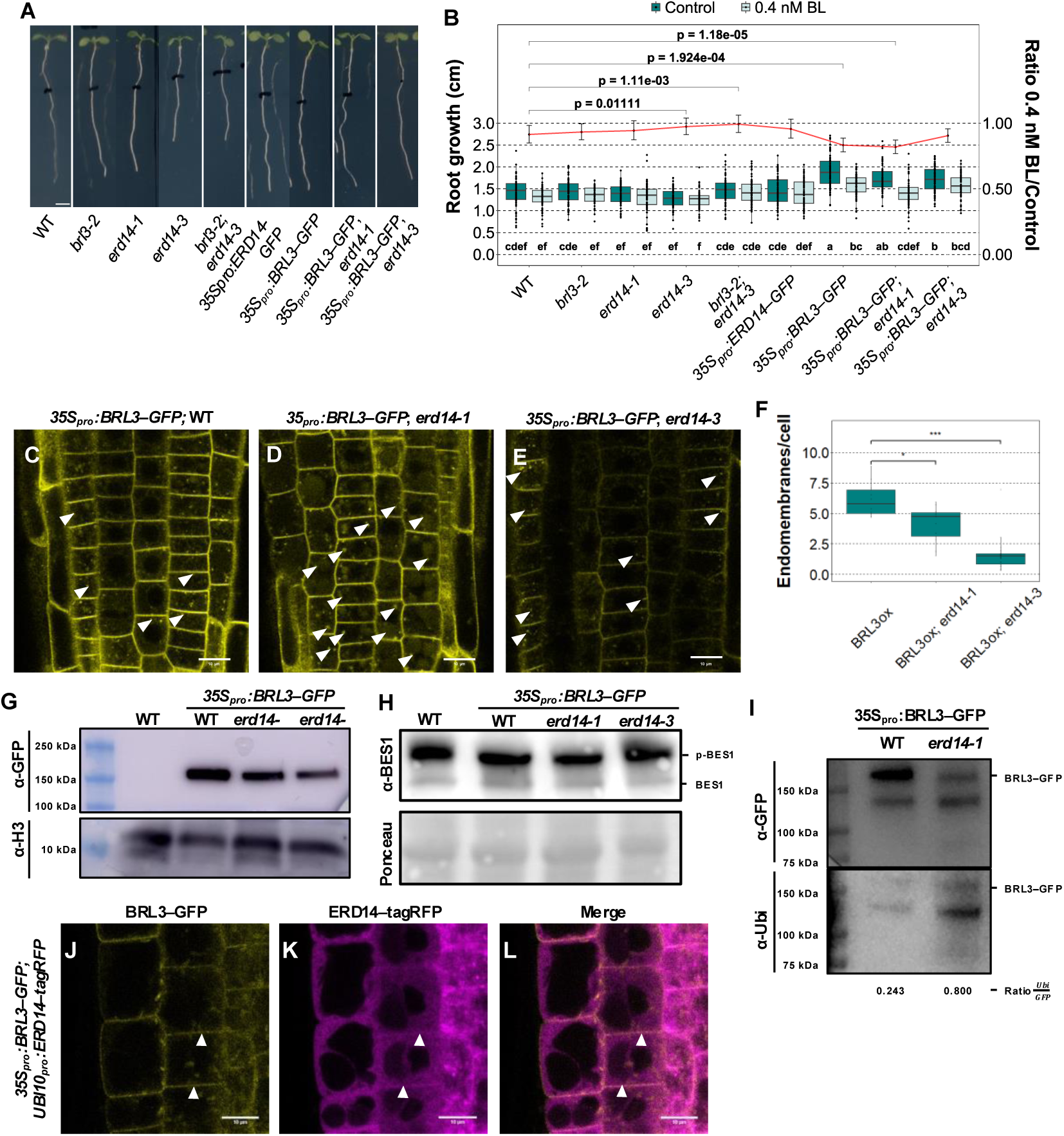
Proper BRL3 function and localization depends on the dehydrin ERD14. **A** Phenotypes of 7-d-old seedlings after 3 d of growth from the black lines that depict the length of the root before treatment. Scale bar: 2.5 mm. **B** Boxplots of root growth (left y axis) after treatment and ratio of root growth 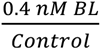 (right y axis) for 7-d-old seedlings grown in control media (½ MS) or ½ MS + 0.4 nM brassinolide. Red line depicts relative root-growth inhibition upon brassinolide treatment (i.e. ratio of BL/control ± CI., n >116 from 3 independent replicates). Letters depict statistically significant differences within each genotype and treatment in a two-way ANOVA plus a Tukey’s HSD test. Statistical significance of the ratios was determined by Welch’s t-test. **C–E** Confocal microscopy of BRL3–GFP subcellular localization in root-epidermal cells. 7-day old transgenic lines express a *35S_pro_:BRL3*–*GFP* cassette in Col-0, *erd14*-*1* and *erd14*-*3* backgrounds, respectively. Arrows indicate BRL3-GFP endomembranes. Scale bar: 10 µm. **F** Quantification of BRL3–GFP endomembranes per cell in *erd14* mutants from epidermal cells. Data are n = 10 different plants. Statistical significance is shown by asterisks after Wilcoxon test (P < 0.05). **G** Western blot of BRL3–GFP abundance by anti-GFP antibody in 5-d-old WT, *35S_pro_:BRL3*–*GFP*, *35S_pro_:BRL3*–*GFP erd14-1* and *35S_pro_:BRL3*–*GFP erd14-3* seedlings. H3 was used as a loading control and probed using anti-H3 antibody. **H** Western blot of BES1 abundance by anti-BES1 antibody in 5-d-old WT, *35S_pro_:BRL3*– *GFP*, *35S_pro_:BRL3*–*GFP erd14-1* and *35S_pro_:BRL3*–*GFP erd14-3* seedlings. For the anti-BES1 blot, the upper and lower bands correspond to the phosphorylated and non-phosphorylated forms of BES1, respectively. Ponceau staining was used as a loading control. **I** Western blot of BRL3–GFP (top) and ubiquitin (Ubi, bottom) abundance with anti-GFP and anti-Ubi antibodies respectively after microsomal enrichment and anti-GFP immunoprecipitation from 7-d-old transgenic lines expressing *35S_pro_:BRL3*–*GFP* in Col-0 and *erd14-1* backgrounds. The ratio is between the mean intensity of bands. **J–L** Confocal microscopy of root-epidermal cells in 7-d-old dual-transgenic lines expressing *35S_pro_:BRL3*–*GFP + UBI10_pro_:ERD14–tagRFP* constructs in the Col-0 background. **I** BRL3–GFP subcellular localization. **J** ERD14*–*tagRFP subcellular localization. **K** Co-localization of BRL3–GFP and ERD14*–*tagRFP at subcellular level. Arrows indicate plasma-membrane localization Scale bar: 10 µm.

Owing to their physical interaction (Fàbregas et al. 2013), we then tested whether ERD14 alters BRL3 stability by introducing a *BRL3* over-expression construct (*35S_pro_:BRL3–GFP*) into *erd14-1* and *erd14-3* knock-down and knock-out mutants, respectively (Supplementary Figure 1). Confocal imaging showed BRL3–GFP fluorescence was lower in *erd14-3* mutant background than a Col-0 background, and the receptor was internalized in endomembranes wherein it is non-functional in both *erd14* mutant backgrounds (Figure 1C–F). Fluorescence reduction was stronger in the *erd14-3* knock-out than *erd14-1*, supporting a model whereby ERD14 contributes to BRL3 stabilization at the plasma membrane. This coincided with lower visible BRL3–GFP internalizations in the erd14 mutants (Figure 1F), hinting at higher BRL3 degradation in absence of ERD14. Western blotting confirmed lower BRL3–GFP abundance in *erd14* mutants than in the Col-0 background (Figure 1G). Moreover, the brassinosteroid signaling pathway was down-regulated in the *erd14* background compared to the WT, inferred by accumulation of phosphorylated BES1 in the *35S_pro_:BRL3–GFP erd14-3* background compared to *35S_pro_:BRL3–GFP* (Figure 1H).

Ubiquitination regulates BRI1 dynamics(Martins et al. 2015). Toward understanding the mechanistic basis of differential BRL3–GFP accumulation on the plasma membrane in *erd14* mutants, we investigated post-translational modifications in *35S_pro_:BRL3–GFP* backgrounds. Using anti-ubiquitin (Ubi) antibodies, we saw that the ratio between Ubi and BRL3–GFP in the microsomal-enriched fraction was higher in the *erd14-1* mutant background than Col-0 (Figure 1I), indicating BRL3–GFP localization on the membrane depends on ubiquitination and that presence of ERD14 in the protein complex helps prevent it.

### BRL3 and ERD14 co-localize at tissue and sub-cellular levels

BRL3 is expressed in the phloem and the quiescent-center cells of the Arabidopsis root apical meristem (Caño-Delgado et al. 2004; Fàbregas et al. 2013). Here we investigated the ERD14–GFP expression pattern in 6-d-old *ERD14_pro_:ERD14–GFP* seedlings by confocal microscopy at higher resolution than before (Nylander et al. 2001). ERD14–GFP was expressed in the columella and the quiescent center of the root tip and was enriched in the root vasculature, coinciding with the known pattern for BRL3–YFP (Caño-Delgado et al. 2004; Salazar-Henao et al. 2016). (Supplementary Figure 2A–D).

At the subcellular level, confocal imaging of stable-transgenic dual over-expression lines showed a well-defined plasma-membrane localization for BRL3–YFP and dual cytosolic and plasma-membrane localization for ERD14–tagRFP (Figure 1J–L). These observations are consistent with transient expression in *Nicotiana benthamiana* (Supplementary Figure 3A–C) and with single stable-transgenic *35S_pro_:BRL3–GFP* and *35S_pro_:ERD14–GFP* Arabidopsis lines (Supplementary Figure 3 G,I). A change in subcellular localization was observed upon osmotic stress induced by 270 mM sorbitol treatment for 30 min, in which BRL3–GFP and ERD14–GFP were released from the plasma membrane and became more cytosolic (Supplementary Figure 3D–F and 3J–L), indicating an internalization pathway for both proteins in response to osmotic stress.

### BRL3 and ERD14 physically interact at the plasma membrane

ERD14 was identified as a BRL3 interactor by LC–MS/MS (Fàbregas et al. 2013). Here, we validated this interaction by co-immunoprecipitation (Co-IP), bimolecular fluorescence complementation (BiFC) and yeast two-hybrid assay (Figure 2A and Supplementary Figures 4–5). For Co-IP experiments, we transiently co-expressed BRL3– GFP and ERD14–3xHA in *N. benthamiana* leaves and pulled down BRL3–GFP using anti-GFP beads, followed by western blot. BRL3–GFP and ERD14–3xHA immunoprecipitated together, confirming they are constituents of the same signalosome complex (Figure 2A). These results were corroborated by BiFC and yeast two-hybrid assays (Supplementary Figures 4, 5A). In both experiments, the BRI1–BKI1 interaction was used as positive control for LRR–receptor-kinase interaction with a disordered protein (so-called unstructured proteins) and BRI1 (Wang and Chory 2006; Jaillais et al. 2011). Otherwise, BRL3 lacks the BKI1-binding domain and BKI1 was not detected in previous mass-spectrometry analysis of BRL3 interactors (Jaillais et al. 2011; Fàbregas et al. 2013; Lozano-Elena and Caño-Delgado 2019), so the BRL3–BKI1 pair was used as negative control. While a BRL3–BKI1 interaction was not detected in our hands, all tested biochemical approaches confirm a BRL3–ERD14 physical interaction (Supplementary Figures 4, 5A).

**Figure 2.**
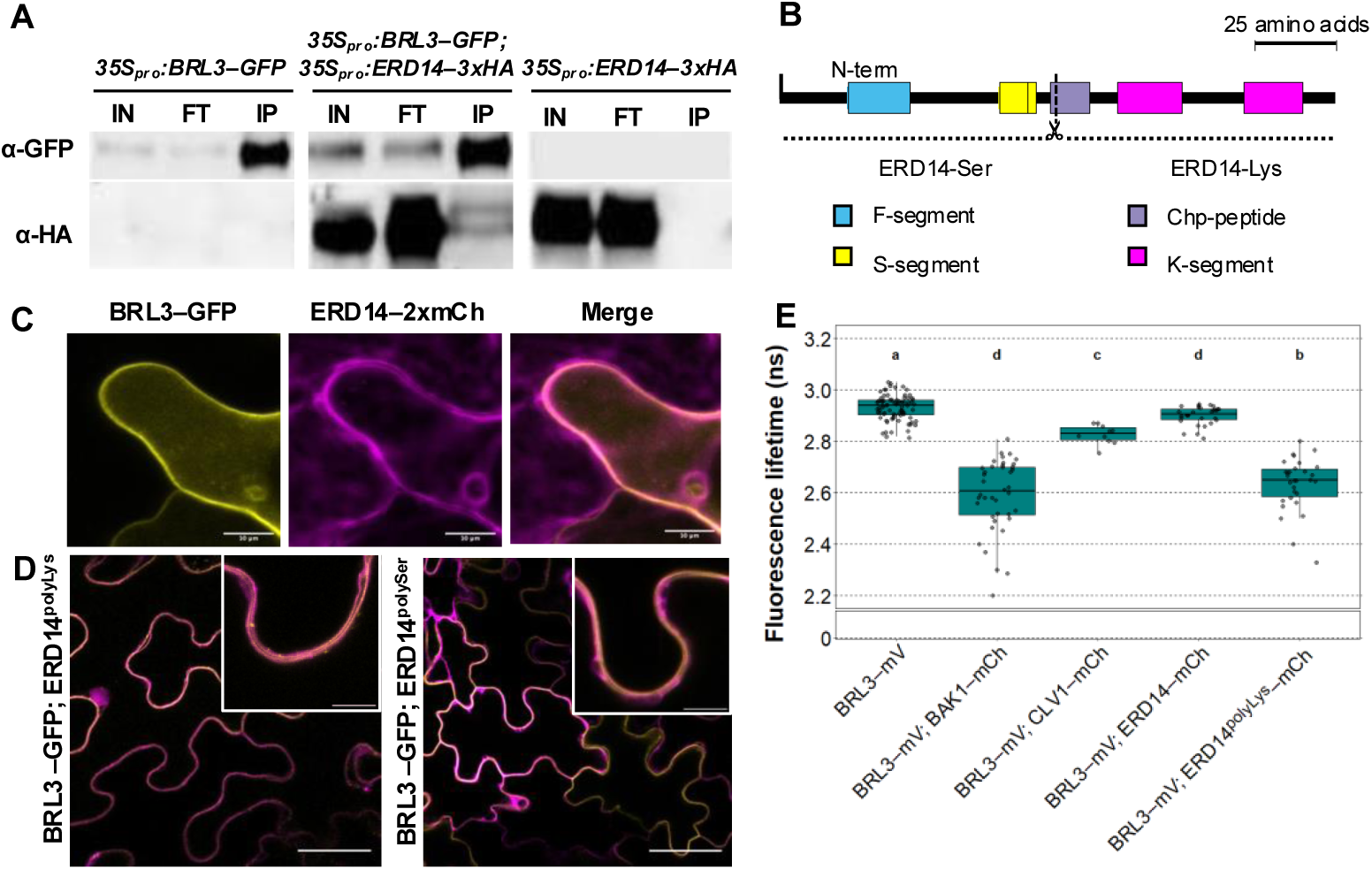
BRL3 physically interacts with ERD14 at the plasma membrane. **A** Co-IP of BRL3–GFP and ERD14*–*3xHA following transient co-expression in *N. benthamiana* leaves. Fractions correspond to the crude extract (input), unbound flow through (FT) and immunoprecipitated using anti-GFP beads (IP). **B** Schematic representation of ERD14 domain organization and the two truncated versions used either side of the dotted line, designated ERD14^polySer^ and ERD14^polyLys^. **C** Confocal microscopy of BRL3–GFP and ERD14*–*2xmCherry transiently co-expressed in *N. benthamiana* leaves. Scale bar: 10 µm. **D** Confocal microscopy of BRL3–GFP with either ERD14^polyLys^*–*2xmCherry or ERD14^polySer^*–*2xmCherry transiently co-expressed in *N. benthamiana* leaves. Scale bar:50 µm. Insets depict the plasma membrane region. Scale bar: 10 µm. **E** FRET–FLIM assay of the physical interaction between BRL3–mVenus (mV) and ERD14–mCherry (mCh). BRL3–BAK1 serves as a positive control and BRL3–CLV1 as a negative control. Data are from n > 10 co-expression events in *N. benthamiana* leaves from three independent replicates. Different letters depict statistically significant difference in fluorescence-lifetimes means determined by one-way Kruskal –Wallis test with Bonferroni correction (*P* < 0.05).

### Different domains of ERD14 regulate its subcellular localization

To study the dynamics and map the interaction of ERD14 with BRL3, ERD14 was split into two truncations (Figure 2B), one containing the serine domain and the F-segment (designated ERD14^Ser^), and the other containing a poly-Lysine (Chp-segment) domain and the two DHN domains (K-segments) that are rich in lysine (designated ERD14^Lys^) (Richard Strimbeck 2017). Confocal imaging of full-length BRL3–GFP and full-length ERD14–2xmCherry showed partial colocalization at the plasma membrane in *N. benthamiana* leaves (Figure 2C). ERD14 truncations showed a well-defined plasma membrane localization for ERD14^polyLys^, whereas ERD14^polySer^ was absent from the plasma membrane but enriched in the cytosol (Figure 2D, Supplementary Figure 4).

To map the interaction, we studied ERD14 truncations in yeast two-hybrid assays (Supplementary Figure 5A) and fluorescence resonance energy transfer measured by fluorescence lifetime imaging (FRET–FLIM, Figure 2E and Supplementary Figure 5B). In yeast two-hybrid, the interaction with ERD14^polyLys^ was stronger than full-length ERD14 (Figure 2E), while the interaction with ERD14^polySer^ could not be addressed due to auto-activation. In FRET–FLIM experiments, the BRL3–BAK1 interaction was used as a positive control (Fàbregas et al. 2013), and the BRL3–CLV1 non-interaction as a negative control. BRL3–mVenus interacted exclusively with ERD14^polyLys^–2xmCherry, which contains the ERD14 chaperone domains (Figure 2E and Supplementary Figure 5B). We tracked the subcellular localization of ERD14 truncations in *N. benthamiana* and saw ERD14^polyLys^–GFP underwent similar localization changes upon stress than full-length ERD14, with the protein being internalized from the plasma membrane region to cytosolic endomembranes (Supplementary Figure 3J–L and Supplementary Figure 6A–C). These results suggest the requirement of the Lys-rich regions of the protein for the localization changes upon osmotic stress. However, ERD14^polySer^ localization or intensity were unchanged (Supplementary Figure 6D–F). Taken together, these results indicate that the Lys domain of ERD14 is essential for the ERD14–BRL3 interaction at the plasma membrane.

### The BRL3–ERD14 heterodimer is required for normal root growth during abiotic-stress adaptation

To address the physiological relevance of the BRL3–ERD14 interaction for the conditional phenotypes of the *brl3-2* mutant and *35S_pro_:BRL3–GFP* seedlings (Fàbregas et al. 2018; Gupta et al. 2023), we examined their performance together with *erd14-3* mutants and *35S_pro_:ERD14–GFP*, under various abiotic-stress conditions. Firstly, we studied the thermo-morphogenic response inferred by comparing hypocotyl growth (Casal and Balasubramanian 2019) at 22°C versus 28°C. The *erd14-3* mutant had reduced hypocotyl elongation (Figure 3A,B) akin to *brl3-2* (Gupta et al. 2023). In contrast to the increased hypocotyl elongation observed in *35S_pro_:BRL3–GFP* plants, *35S_pro_:ERD14–GFP* plants had an attenuated response (Figure 3A,B). A similar reduction is observed in the *35S_pro_:BRL3–GFP erd14-3* line that reverted *35S_pro_:BRL3–GFP* hyper-elongation to WT levels (Figure 3A,B). This suggests that *ERD14* is required for *BRL3*-mediated thermo-morphogenesis in the hypocotyl.

**Figure 3.**
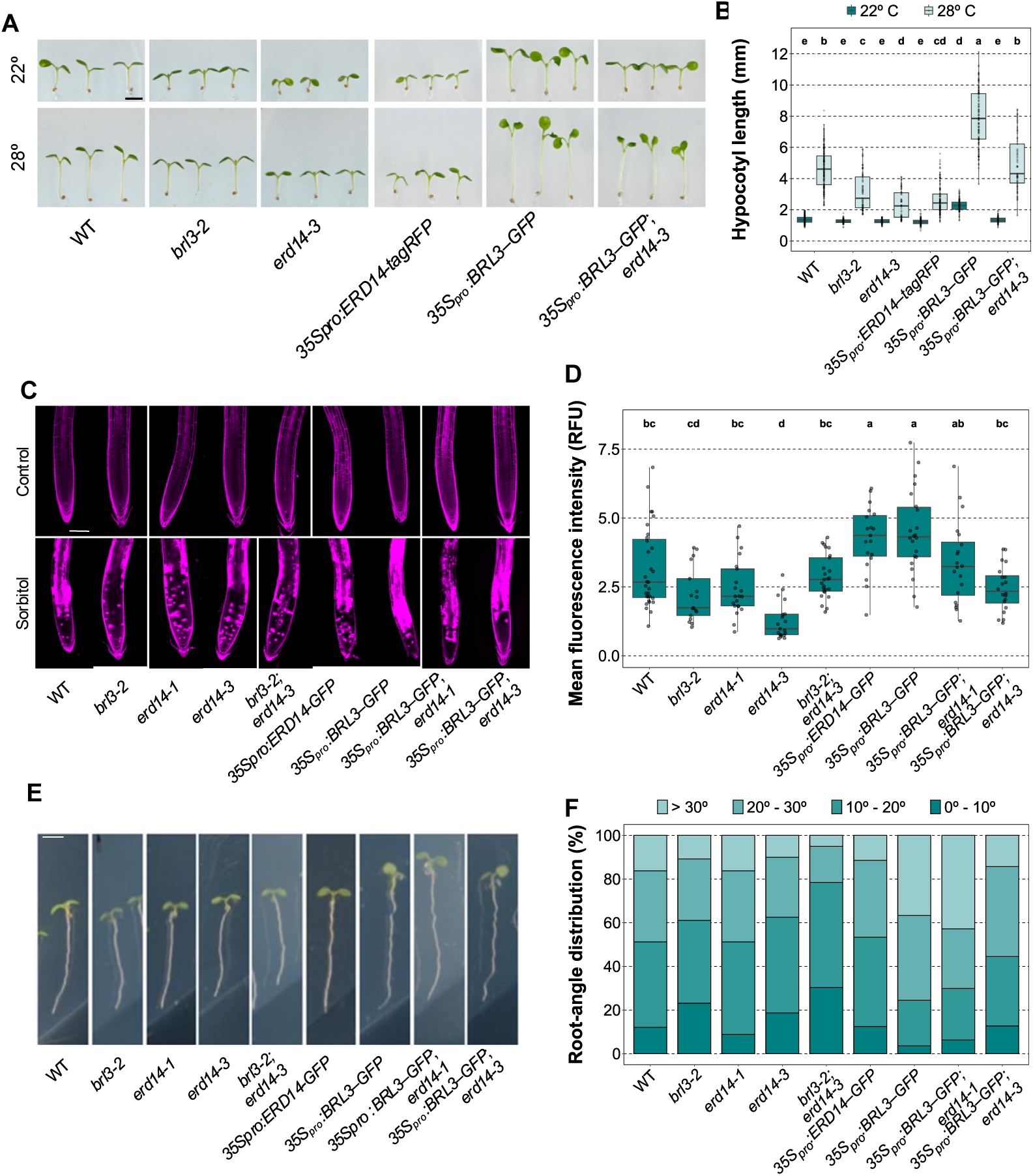
BRL3–ERD14 signaling is active during abiotic-stress adaptation. **A** Hypocotyl phenotypes after 6 d of continuous growth at 22°C (left) or 28°C (right). Scale bar: 1 mm. **B** Quantification of hypocotyl length after 6 d at 22°C or 28°C. Data are n > 60 hypocotyls from three independent replicates. The effects of genotype and growth temperature on the means of hypocotyl length were determined by two-way ANOVA and Tukey’s HSD test. Different letters depict statistically significant differences (*P* < 0.05). **C** Confocal microscopy of 6-d-old roots stained with propidium iodide (false-colored magenta) after 24 h in control media (1/2 MS, top) or 270 mM sorbitol (bottom). Scale bar: 10 µM. **D** Propidium-iodide intensity as a proxy (relative fluorescence units / RFU) for cell death. Data are n > 17 root meristems from two independent replicates. Different letters depict statistically significant differences in means determined by one-way Kruskal–Wallis test with Bonferroni correction (*P* < 0.05). **E** Root-curvature phenotypes for 7-d-old seedlings undergoing a hydrotropic response after exposure to 270 mM sorbitol for 24 h. Scale bar: 5mm. **F** Discrete distribution of root-curvature angles in different intervals for plants grown as in E. Data are n > 79 roots from 10 independent replicates.

Secondly, using propidium iodide staining and confocal microscopy, we measured cellular uptake of the dye and quantify programmed cell death caused by osmotic stress for 24 h with 270 mM sorbitol (Duan et al. 2010). *erd14-1, erd14-3, brl3-2* and *brl3-2 erd14-3* plants all had fewer damaged cells than Col-0 (Figure 3C,D), whereas *35S_pro_:BRL3–GFP* had more dead cells, in agreement with previous results (Fàbregas et al. 2018). *35S_pro_:BRL3–GFP erd14*-*3* plants exhibited a WT-like phenotype, while the presence of the *erd14* mutation reverted the *35S_pro_:BRL3–GFP* phenotype (Figure 3CD). Lastly, we tested the capacity of roots to detect and avoid osmotic stress by bending towards water-rich media in hydrotropism assays (Figure 3E) (Takahashi et al. 2002). The *erd14-1*, *erd14-3, brl3-2* and *brl3-2 erd14-3* mutants showed an impaired hydrotropic response compared to Col-0 and *35S_pro_:BRL3–GFP* plants, in which this response was enhanced (Figure 3F and Supplementary Figure 7) (Fàbregas et al. 2018). In contrast, while no statistically significant differences were observed between *35S_pro_:ERD14–GFP* and Col-0 plants (Figure 3F), root bending for the *35S_pro_:BRL3–GFP erd14*-*3* plants was reduced to WT-level. Taken together, our data support the notion that ERD14 may act as a partner for BRLs, important for full BRL3 activity to ensure proper root adaptation to abiotic stress.

### The BRL3–ERD14 interaction is essential to confer drought resistance in mature plants

To further understand the physiological relevance of this interaction, we studied the performance of mature plants during drought-stress adaptation. Known drought resistance of *35S_pro_:BRL3–GFP* plants (Fàbregas et al. 2018) was reverted to the WT situation in adult *35S_pro_:BRL3–GFP erd14*-*3* plants (Figure 4A,B). Additionally, *brl3-2* was more sensitive to drought when compared to WT, while *erd14-1* was slightly more resistant (Figure 4A,B, Supplementary Figure 8). We also analyzed water usage by scoring time taken for plants to reach 10% of field capacity (i.e. 90% of the available water was consumed), as per done in Fàbregas et al., 2018. Consistent with reduced survival after drought, *35S_pro_:BRL3–GFP erd14*-*3* plants did not have reduced water usage, as seen for *35S_pro_:BRL3–GFP* plants (Figure 4C). These results were also consistent with enhanced BES1 de-phosphorylation in *35S_pro_:BRL3–GFP* plants and its reversion to WT levels in *35S_pro_:BRL3–GFP erd14*-*3* (Figure 1H). This finding suggests that the activation of brassinosteroid signaling through BRL3 is required to promote drought resistance and that this activation is dependent on ERD14 chaperone function.

**Figure 4.**
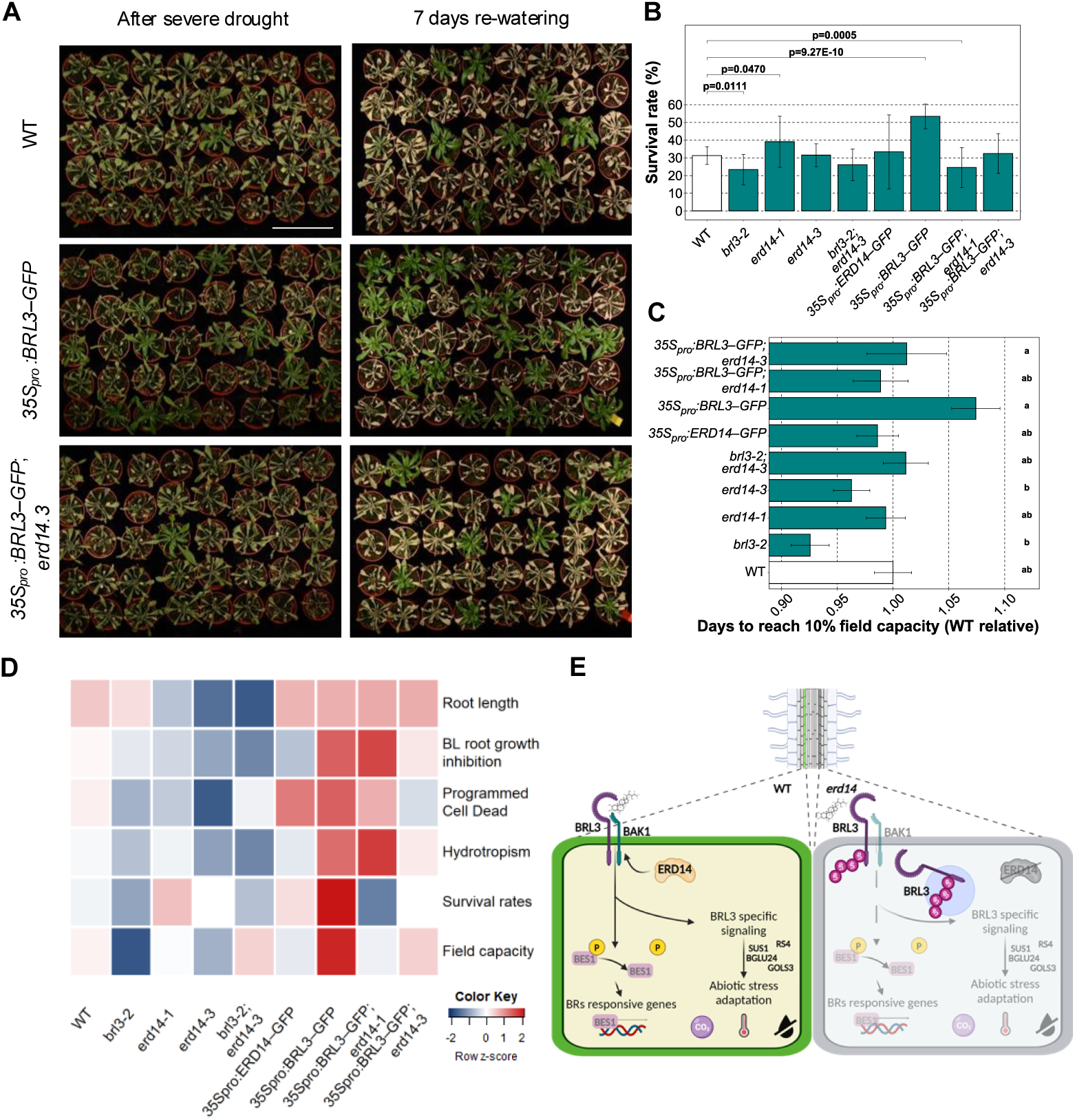
The BRL3–ERD14 signalosome accounts for multiple drought-adaptation traits. **A** Phenotypes of 40-d-old plants after severe drought stress (left) and following 7 d of re-watering (right). Scale bar: 10 cm. **B** Plant survival after 7 d of re-watering for plants shown in A. Averages from n > 111 plants from at least three independent biological replicas. Bars are the mean ± S.E.. Statistical difference shown by p-values from a chi-squared test for survival ratios compared to Col-0. **C** Time taken to reach 10% field-capacity. Bar plots depict the normalized time in comparison to WT needed to reach 10% of the soil field capacity. Data are n > 58 plants studied across three independent replicates. Different letters represent statistically significant differences in one-way Kruskal–Wallis test with Bonferroni correction (*P* < 0.05). Bars represent the standard deviation. **D** Trait matrix summarizing relative responses amongst genotypes for all physiological experiments performed. The colored bar depicts relative values for scaled data. **E** Proposed molecular model of ERD14 function in BRL3-dependent signaling. In the wild type, ERD14 physically interacts with and contributes to BRL3 localization at the plasma membrane in vascular tissues, inducing canonical brassinosteroid signaling to consequently regulate expression of osmoprotectant-biosynthesis genes and confer drought resistance. In *erd14* mutants, BRL3 is ubiquitinated and subsequently internalized within endomembranes. Brassinosteroid signaling is down-regulated and osmoprotectant-biosynthesis genes are not induced, resulting in plants sensitive to drought stress.

Drought resistance of *35S_pro_:BRL3–GFP* plants is linked to over-accumulation of osmoprotectant metabolites and up-regulation of genes involved in their biosynthesis (Fàbregas et al. 2018). The expression of *BGLU24*, *GOLS3*, *RS4*, *SUS1*, *TTPB* and *TTPH* was restored to WT levels in *35S_pro_:BRL3–GFP erd14*-*3* plants (Supplementary Figure 9), indicating that ERD14 is crucial to establish the osmoprotectant signature of these plants (Fàbregas et al. 2018).

## Discussion

The characterization of mechanisms controlling brassinosteroid signaling at the receptor level has become a key point for understanding the role of this hormone towards plant adaptation to adverse environments (Lozano-Elena and Caño-Delgado 2019; Gupta et al. 2020; Nolan et al. 2020). In this study, we delved into the molecular mechanisms by which the BRL3 receptor promotes drought resistance. We showed that the dehydrin ERD14 interacts with and chaperones BRL3 to the plasma membrane, wherein the receptor activates brassinosteroid signaling in response to environmental stress, to induce osmoprotectant biosynthesis genes and promote drought resistance (Figure 4D,E). We have demonstrated that the interaction of BRL3 with ERD14 prevents the ubiquitination of the receptor, avoiding its degradation and increasing its levels at the plasma membrane.

Ubiquitination is a well-known mechanism regulating signaling pathways upon biotic and abiotic stress (Lu et al. 2011; Trujillo 2021; Singh et al. 2022). Moreover, ubiquitination regulates brassinosteroid signaling via modification of the BRI1 receptor, whereby it affects plant growth and development (Martins et al. 2015). It is therefore conceivable that ERD14 is responsible for BRL3 localization upon stress by preventing receptor ubiquitination. Further supporting our hypothesis, the brassinosteroid-mediated action of BRL3 and its downstream effector BES1 depend on ERD14 expression, thus confirming that ectopic signaling induces plant adaptation to drought. Future structural studies of the interaction between BRL3 and ERD14 will provide relevant information about its mode of action that could be used to modulate downstream signaling. However, due to the intrinsically disordered nature of ERD14, the structural characterization of this interaction will be technically challenging even for the new prediction tools such as AlphaFold2 (Jumper et al. 2021), that was unable to provide an interaction model for BRL3-ERD14 interaction with enough confidence when tested. Despite this, our results evidence that the regions of ERD14 containing the K-segments and the ChP-segment are responsible for the interaction with BRL3. This is consistent with previous studies on ERD14 chaperone activity *in vitro* indicating the K-segments are the most relevant domains for this (Murvai et al. 2021). Moreover, K-segments are responsible for conferring heat-stress resistance in *E. coli* (Murvai et al. 2021).

Following the model proposed by Murvai et al., 2021, ERD14 may structurally favor the simultaneous association of BRL3 with several partners form the BRL3 signaling complex at the membrane to chaperone this interaction under abiotic stress and thus allowing proper activation of the BRL3 signalosome (Murvai et al. 2021). Introduction of *erd14* restores *35S_pro_:BRL3–GFP* phenotypes to WT levels, thereby indicating that no other protein can completely compensate ERD14 function in the BRL3 signaling pathway in these conditions. At the subcellular level, *erd14* results in aberrant BRL3*–*GFP internalization and BRL3*–*GFP abundance is lower. A similar effect is known for other chaperones of receptor-like kinases, such as LORELEI-LIKE GLYCOSYL PHOSPHATIDYL INOSITOL-ANCHORED PROTEIN 1 (LLG1), which delivers FERONIA (FER) to the plasma membrane by interacting with its extracellular domain (Li et al. 2015). In the *llg1-2* mutant, FER remains in the endoplasmic reticulum. DE-ETIOLATION IN THE DARK AND YELLOWING IN THE LIGHT (DAY) is a BRI1 chaperone that interacts with its extracellular domain and is responsible for plasma-membrane localization (Li et al. 2015; Lee et al. 2021).

This study provides a new perspective for coping with a changing environment in which plants will imminently face adverse conditions such as elevated temperatures or drought. BRL3 has been identified in several conditional phenotypes associated with environmental adaptation and abiotic-stress responses (Gupta et al. 2023) and its overexpression in Arabidopsis results in enhanced adaptation to select abiotic stressors (Fàbregas et al. 2018). By modulating the activity of the BRL3 complex through genome editing and bioengineering, as showed by ERD14 affecting BRL3 ubiquitination, we can potentially deploy a tool for generating plants better suited to adverse environmental conditions. Future studies of the BRL3 interactome and signalosome are expected to lead to a deeper knowledge of BRL3 dependent adaptation to abiotic stress and the potential translational use for enhancing drought-resistance traits in crops without compromising overall plant productivity and yield.

## Materials and methods

### Growth conditions and plant material

*Arabidopsis thaliana* Columbia-0 was used as the wild type (WT) with all mutants and transgenic lines generated in this background, namely *erd14-1*, *erd14-3*, *brl3-2*, *brl3-2 erd14-3*, *35S_pro_:BRL3–GFP*, *BRL3_pro_:BRL3–YFP, 35S_pro_:ERD14–GFP, UBI10_pro_:ERD14–tagRFP, ERD14_pro_:ERD14–GFP*, *35S_pro_:BRL3–GFP erd14-1*, *35S_pro_:BRL3–GFP erd14-3* and *35S_pro_:BRL3–GFP UBI10_pro_:ERD14–tagRFP*. Additional information can be found in Supplementary Table 1.

Seeds were surface-sterilized in 35% v/v sodium hypochlorite for 5 min with agitation followed by 5x 5-min washes in sterile H_2_O. Seeds were stratified at 4°C for 48*–* 72 h and germinated on plates containing sterile half-strength Murashige and Skoog medium (1/2 MS) without sucrose (Murashige and Skoog 1962). Plants were grown under long-day photoperiod with 16 h illumination at 150 μmol photons m^−2^ s^−1^ and 8 h darkness, all at 22–23°C. All lines were genotyped by PCR before experimentation and transgenic lines were selected according to their selectable marker (Supplementary Table 1). Progenitor lines were grown on soil under long-day conditions (different mixtures of autoclaved soil supplemented with perlite and vermiculite for aeration were used). All seeds collected were left to dry for 3–4 d prior to use.

*Nicotiana benthamiana* plants with the *rdr6i* genotype (Schwach et al. 2005) were grown for 2–3 weeks in greenhouses under natural illumination conditions in autoclaved soil supplemented with perlite and vermiculite for aeration. Temperature was maintained between 24–28 °C in the day and night, and relative humidity was between 50–65%. Plants were directly germinated on soil.

### ERD14 cloning and generation of transgenic lines

*ERD14* (AT1G76180) sequences were amplified from genomic DNA by PCR (Supplementary Table 2) to produce a 642-bp coding-region fragment without codon stop but including the single 87-bp intron, or from cDNA, avoiding the 87-bp intron. The *ERD14* promoter was amplified from genomic DNA, containing the complete intergenic region and 2,667bp upstream of the *ERD14* translational start codon (i.e. the promoter region). *ERD14* and the *ERD14* promoter were subcloned into pDONR 221 and pDONR P4P1r entry vectors respectively using Gateway system (Invitrogen). To generate the expression constructs *ERD14_pro_:GUS–GFP*, *ERD14_pro_:ERD14–GFP*, *35S_pro_:ERD14–3xHA*, and *35S_pro_:ERD14–GFP*, the *ERD14* coding sequence and promoter region were introduced into binary vectors R4L1pGWB632, R4pGWB604, pGWB414 (Nakagawa et al. 2007) and pH7FWG2 (Karimi et al. 2002) respectively. Binary vectors except the one containing *35S_pro_:ERD14–3xHA* were transformed into *Agrobacterium tumefaciens* GV3101 and positive colonies were used to transform Arabidopsis by floral dipping.

Selection of transgenic Arabidopsis was done using specific antibodies indicated in the Supp. Table 1. At least 10 T1 lines were selected, and they were screened for single insertion segregation in T2. Three of them with consistent phenotype and expression were carried until homozygous T3. Chi-square testing was performed to validate the proportions of 75:25 in T2 and 100:0 in T3 (resistant:susceptible). Expression was assayed via fluorescence or histochemistry in case of GUS. The overexpression of ERD14 was also tested by RT–qPCR with the primes listed in Supplementary Table 3.

### Brassinolide-sensitivity assays

Brassinolide treatments were carried out to examine root-growth inhibition on vertical plates containing 0.4 nM brassinolide (Wako, Japan) over a period of 6 d. Brassinolide was serially diluted from a 10 mM stock dissolved in DMSO. Images were captured daily with NIKON D7000 and NIKON Z50 cameras and root measurements were quantified using Image J (https://imagej.net/software/fiji/). Average values for root length were calculated for each genotype.

### Hydrotropism assays

Seedlings were germinated in solid ½ MS without sucrose plates and grown in the vertical position for 6 d. Then, the lower part of another solid ½ MS without sucrose plate was removed and ½ MS with 270 mM sorbitol + agar was added to simulate a situation of reduced water availability. The seedlings were then transferred to the sorbitol-containing plates with the tip 5 mm away from the sorbitol-media border at the diagonal axis to allow diffusion and avoid direct contact with sorbitol media. The new plates were placed on a 45° angle during 24 h to scape gravitropism effect. Photographs were captured with NIKON D7000 and NIKON Z50 cameras and root-angle measures were determined using Image J (http://rsb.info.nih.gov/ij/). Average values for bending angle from the transference point were measured for each genotype.

### Drought simulation field-capacity assays

Ten-day-old seedlings were grown on ½ MS agar plates and transferred individually to pots containing 30 ± 1 g of substrate (11:1:0.5 v/v/v ratio of Substrate 2 (Klasmann-Deilmann):vermiculite:perlite). For each survival experiment, plants were grown in well-watered conditions until 3 weeks old before being subjected to severe drought stress by withholding water for 10 d, followed by re-watering. After a 7-d recovery period with regular watering, the surviving plants were photographed, counted, and Chi-squared testing carried out for survival ratios compared to Col-0 (p-value < 0.05). The surviving plants were counted as such if they still presented green aerial tissues and were able to keep growing, as done in Fàbregas et al. (2018).

For field-capacity assays, plants were grown in well-watered conditions until 3 weeks old before being subjected to severe drought stress by withholding water for the number of days required to reduce the field capacity of the pots to 10%. The field capacity corresponds to the amount of water available in the pot and it was calculated by measuring the weight difference between the saturated media at 100% field capacity (0% of water loss) and the sample. Weight corresponding to the substrate and plastic pot (0% field capacity) was deduced from the measuring using fully dried samples (100% of water loss). Plant weight was not taken into consideration for the calculations. The time required to arrive to 10% of field capacity was normalized within replicates for each genotype using Col-0.

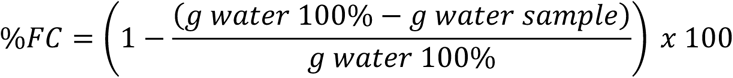

### Cell-death assays

Cell-death assays were performed with 5-d-old seedlings grown on ½ MS agar plates and then transferred to ½ MS agar plates containing 270 mM sorbitol for 24 h. Root tips were then visualized by confocal microscopy (detailed below). Image areas showing evidence of staining with propidium iodide were quantified in a window of 400 µm^2^ and analyzed using Image J (https://imagej.net/software/fiji/). Relative fluorescence units (RFU) of the stressed samples were normalized using non stressed samples within the same replica to reduce batch-effect differences between days due to external variables of propidium-iodide stocks and visualization parameters.

### Western blotting

Total proteins were extracted from 1 g of frozen seedling tissue in 1 mL of extraction buffer (50 mM Tris-HCl pH 7.5, 150 mM NaCl, 1% v/v Triton X-100) supplemented with 1x Plant Protease Inhibitor Cocktail (P9599 Sigma) and Phosphatase Inhibitor Cocktail 3 (P0044 Sigma) on ice. Extracts were vortexed for 1 min then centrifuged in a tabletop microfuge for 10 min at 5,000 g at 4°C to remove cellular debris. Microsomal enrichment for ubiquitination assay was performed as described in Martins et al. (2017). Aliquots of 40 µL were taken and 1x Laemmli loading buffer (60 mM Tris-Cl pH 6.8, 2% w/v SDS, 10% v/v glycerol, 5% v/v β-mercaptoethanol, 0.01% w/v bromophenol blue) was added and samples were denatured at 95°C for 5 min.

Mini-PROTEAN TGX Stain-Free precast gels for polyacrylamide gel electrophoresis (SDS–PAGE) were used and equal volume of denatured protein was loaded for all samples. Protein samples were run at 100 V for 15 min then 200 V for 25 min, before transfer to a PVDF membrane (Millipore) by blotting for 1 h at 100 V at 4°C under agitation. Proteins on the membrane were stained using Ponceau Red solution (Sigma) and photographed with a Chemidoc-touch (BioRad). Membranes were blocked using 5% w/v milk dissolved in 1x TBST for 2 h at room temperature. Primary antibodies were diluted in 1% w/v milk dissolved in 1x TBST with 1/1,000 anti-BES1 (Yu et al. 2011), 1/2,000 anti-GFP (Living Colors), 1/5000 anti-histone 3 (NEB) and 1/5,000 anti-ubiquitin (P4D1-A11, Merck). Secondary antibodies were donkey anti-rabbit IgG (AP182P Sigma-Aldrich) for BES1 and histone 3 and donkey anti-mouse IgG (AP192P Sigma-Aldrich) for GFP and ubiquitin. Both were diluted at 1/10,000 in 1% w/v milk dissolved in 1x TBST. SuperSignal™ West Femto Maximum Sensitivity Substrate (Thermo Scientific) was used as substrate for the revealing reaction. Imaging was performed with ImageQuant 800 western-blot imaging system (Amersham).

### Co-immunoprecipitation assay

Co-immunoprecipitation experiments were done as described by Martins et al. (2017) using the μMACS GFP Isolation Kit (Miltenyi Biotec), following the extraction procedure indicated above.

### Yeast two-hybrid assay

Yeast two hybrid assays were performed using pGBKT7-GW and pGADT7-GW Gateway vectors from the GAL4 Two-Hybrid System (Clontech) (Chini et al. 2009). *BRI1* (kinase domain amino acids 814–1,196), *BRL3* (kinase domain amino acids 797–1,164), *ERD14* (either as full length or truncations of 1–90 aa and 91–185 aa) and *BKI1* (full length) were introduced into expression vectors by LR Clonase recombination. Vectors were co-transformed into *Saccharomyces cerevisiae* YM4271a and PJ694α by lithium-acetate transformation (Gietz and Schiestl 2007; Gallemí et al. 2017). Selection for reporter activation was performed in SD–LWH plates. After 3 d of incubation at 30°C, colonies were photographed with NIKON D7000 and NIKON Z50 cameras.

### Bimolecular fluorescence-complementation assay

The coding sequences of *BRI1*, *BRL3*, *ERD14* (either as full length or truncations of 1–90 aa and 91–185 aa) and *BKI1* were inserted by LR Clonase recombination into pBiFC binary expression vectors pGTQL1221YC and pGTQL1211YN (Lu et al. 2010). Plasmids were transformed into *A. tumefaciens* GV3101 by electroporation and infiltrated into 2–3-week-old *N. benthamiana* leaves as described in Waadt and Kudla (2008). Three to five days after infiltration, leaves were observed under an Olympus FV1000 inverted confocal microscope (excitation wavelength 515 nm, emission wavelength 527 nm, BF position 530 nm, BF range 100 nm).

### Confocal-microscopy imaging

Confocal images were captured with a FV 1000 Olympus confocal microscope after propidium iodide staining (PI, 10 µg/mL) by mounting the samples on PI solution immediately before visualization. PI and GFP were detected with a bandpass 570–670 nm (572 nm emission) filter and 500–545 nm (510 nm emission) filter after 559 nm and 488 nm excitation, respectively. Images were taken either in the middle plane or the in the epidermal layer of 6-d-old roots. Fluorescence intensity was quantified with ImageJ using the Integrated Density value obtained from individual plants.

Short sorbitol treatments were performed in liquid ½ MS for 20 min and directly observed by confocal microscopy. Plant-cell-death experiments were performed, imaged and quantified as per Fàbregas et al. (2018). Images in Figure 1 (J–L) and Supplementary Figure 2 were captured using a Confocal Zeiss LSM980 - Elyra 7 system, detecting YFP, GFP, PI and tagRFP with excitation wavelength 514 nm, 488 nm, 514 nm and 558nm respectively and emission wavelength 524 nm, 509 nm, 617 nm and 583 nm using a bandpass 490–552 nm filter, 490–552 nm filter, 579–693 nm filter and 561–693 nm, respectively.

### Fluorescence lifetime imaging microscopy (FLIM)

Coding sequences of *BRL3, CLV1, ERD14* and *ERD14-Lys* without translational stop codons were used for FLIM measurements in *N. benthamiana.* These fragments were obtained by PCR and cloned into pABindmVenus and pABindmCherry Gateway vectors (Berleth et al. 2019) containing the β-estradiol-inducible promoter and a C-terminal mVenus or mCherry reporter fusion, by LR reaction (Invitrogen). *Agrobacterium tumefaciens* GV3101 harboring the p19 silencing-suppressor plasmid was transformed and used for transient transformation of *N. benthamiana* epidermal leaf cells. After 72 – 96 h, transient expression and visualization of mVenus- or mCherry-tagged fusion proteins was induced with 20 µM β-estradiol was carried out as described in Berleth et al. (2019).

### FLIM data acquisition and analysis

FLIM measurements were carried out with a confocal laser-scanning microscope (inverted LSM780, Zeiss) equipped with an additional time-correlated single-photon counting device with picosecond time resolution (Hydra Harp 400, PicoQuant). Excitation of mVenus was performed with a pulsed (32 MHz) diode laser at 485 nm and 1 µW at the objective (40x water immersion, C-Apochromat, NA 1.2 Zeiss). Emission was detected at the same objective and detected with SPAD detectors (PicoQuant) using a narrow-range bandpass filter (534/35, AHF). A series of 40 frames was acquired with each image taken at 12.5 µs pixel dwell time and a resolution of 138 nm/pixel in a 256 x 256 pixel image.

For analysis, 40 frames were merged into one image and analyzed using the SymPhoTime ‘GROUPED FLIM’ tool (PicoQuant). Prior to analysis, a region of interest (ROI) with a threshold of 130 counts was set to exclude background signal from chloroplasts. If necessary, chloroplasts were excluded manually from the ROI. A histogram of fluorescence decay was built from all photons of the ROI. For donor-only samples, a mono-exponential fit model was used to calculate fluorescence lifetime for all photons for the ROI, while for FRET samples containing mVenus and mCherry, a bi-exponential fit model was used. For reconvolution in the fitting process, potassium-iodide-quenched erythrosine was used to measure instrument-response function (Weidtkamp-Peters and Stahl 2017). FLIM images were created by analyzing the fluorescence lifetime of photons from each individual pixel of a merged image with SymPhoTime (PicoQuant). Individual pixels are color-coded according to their fluorescence lifetime.

### RNA isolation, cDNA synthesis and RT–qPCR

Root-tissue samples from 6-d-old plants were harvested and snap-frozen with liquid nitrogen then homogenized using a TissueLyser (QIAGEN). RNA was purified by Maxwell RSC Plant RNA Kit (Promega) following the manufacturer’s instructions and concentration was determined with NanoDrop-1000 spectrophotometer (Thermo Scientific). RNA quality was determined by agarose gel electrophoresis and inspection of ribosomal RNA banding patterns. Reverse transcription was performed using NZY-First Strand cDNA synthesis kit (NZYTech) following manufacturer’s instructions with 1 μg of RNA.

RT-qPCRs were run in triplicate from 5 ng of cDNA in LightCycler 480 SYBR Green I master mix (Roche) in a LightCycler 480 instrument (Roche) using 96-well plates according to the manufacturer’s recommendations. *Actin2* (AT3G18780) (Betegón-Putze et al. 2021) was used as reference gene for normalizing relative expression. Relative expression was calculated according to (Schmittgen and Livak 2008). Primers used are indicated in Supplementary Table 3. To assess PCR reaction efficiencies, standard curves were created using serial dilutions of cDNA. Linear regression between the amount of cDNA template (log [10]) and the cycle threshold (C_T_) value was plotted. Primers with R^2^ > 0.97 were deemed efficient.

### Statistical analysis

All plots, data treatment and statistical analysis were performed using R 4.3.1 (https://www.r-project.org/) and Microsoft Excel v2407 (http://microsoft.com).

After data collection, Saphiro–Wilk testing was done to determine whether data were normally distributed or not (Shapiro and Wilk 1965). This determined the choice of a non-parametric analysis for ratio comparison (Figure 1B), FRET–FLIM (Figure 2E), sorbitol-induced programmed cell death (Figure 3D), hydrotropism (Supplementary Figure 7), drought (Figure 4B) field capacity (Figure 4C), RT–qPCRs (Supplementary Figures 1 and 9), endomembrane quantification (Figure 1F, Supplementary Figure 3I and Supplementary Figure 6C) and fluorescence intensity quantification (Supplementary Figure 3L and Supplementary Figure 6F).

Non-parametric Welch’s t-test (Ruxton 2006) or Wilcoxon test (Wilcoxon 1945) was used for paired comparisons, whereas one-way Kruskal–Wallis plus Bonferroni correction testing was applied for multiple analysis (Kruskal and Wallis 1952; Haynes 2013).

For the drought assays, frequency chi-squared test for non-parametric data was used (Pearson 1992), assuming p < 0.05 to indicate statistically significant differences.

Root-length (Figure 1B) and hypocotyl-elongation (Figure 2B) statistics were analyzed by parametric tests, such as two-way ANOVA plus Tukey’s HSD correction (Tukey 1949; Kaufmann and Schering 2014). Some of the samples of these assays were not perfectly fitting a normal distribution according to Saphiro–Wilk testing by minor differences in the p value, but for simplifying the two-way analysis data were assumed to do so.

## Acknowledgments

We thank Dr Y. Yin for providing anti-AtBES1 antibody. A.I.C.-D. has received funding from the European Research Council (ERC) under the European Union’s Horizon 2020 research and innovation program (grant agreement 683163). A.I.C.-D. is a recipient of a BIO2016-78150-P and BIO2020-118218RB-100 grants funded by the Spanish Ministry of Economy and Competitiveness and Agencia Estatal de Investigación (MINECO/AEI) and Fondo Europeo de Desarrollo Regional (FEDER). A.G. has received funding by a postdoctoral fellowship from the Severo Ochoa Programme for Centers of Excellence in R&D 2016–2019 from the Ministerio de Ciencia e Innovación (SEV-2015-0533). F.L.-E and N.F. received funding from FEDER-BIO2016-78150-P and PIRSES-GA-2013-612583. I.H.G, A.G., F.L.-E., M.M.-B, M.K., N.F. and D.B. are funded by ERC2015-CoG–683163 granted to A.I.C.-D. I.H.G is funded by grant FPU19/04332 from the Spanish Ministry of Universities. S.P. received a contrato Torres Quevedo (PTQ) 2022 pre-doctoral funding (DIN2022-012875), funded by MCIU and AEI. We acknowledge financial support from Spanish Ministry of Science and Innovation-State Research Agency (AEI) through the ‘Severo Ochoa Programme for Centres of Excellence in R&D’ SEV-2015-0533 and CEX2019-000902-S and support from the CERCA Programme/Generalitat de Catalunya. S.A. and A.R.F. acknowledge funding for the PlantaSYST project by the European Union’s Horizon 2020 Research and Innovation Programme (SGA-CSA no. 664621 and no. 739582 under FPA no. 664620). The funders had no role in study design, data collection and analysis, decision to publish, or preparation of the manuscript.

## Author contributions

Conceptualization: ACD, MMB

Methodology: IHG, MMB, SP, AG, FLE, DB, MK, NF, VIS, YS

Investigation: MMB, IHG, ACD, SP, NF

Visualization: AG, FLE, MM, IHG

Funding acquisition: ACD, IHG

Project administration: ACD

Supervision: ACD

Writing – original draft: ACD, MMB, IHG

Writing – review & editing: ACD, MMB, IHG, SP, FLE, AG, NF, MK, VIS, YS

## Data and materials availability

All data are available in the main text or the supplementary materials.

## Supplementary figure legends

**Supplementary Figure S1.**
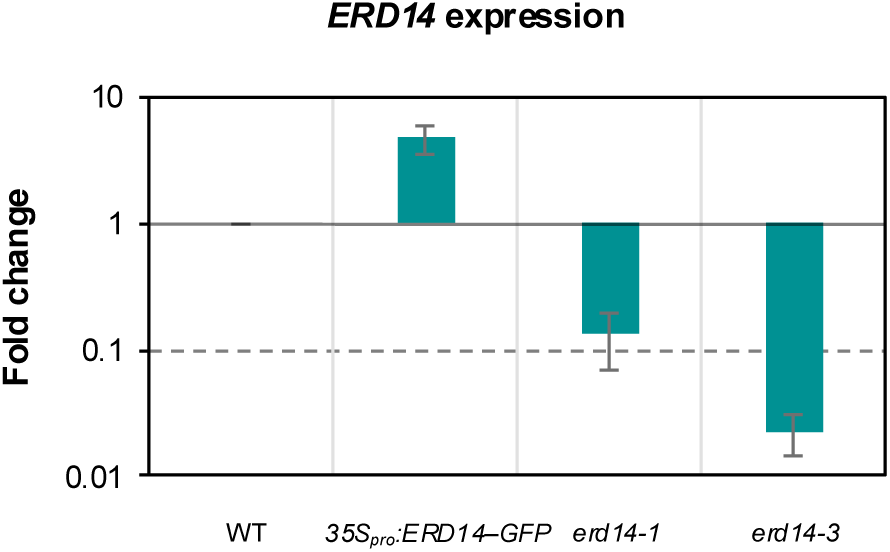
Analysis of *ERD14* expression in transgenic *35S_pro_:ERD14–GFP* and insertion*-*mutant lines. RT–qPCR analyses of *ERD14* expression in *35Spro:ERD14*–*GFP*, *erd14-1*, and *erd14-3* lines relative to *ERD14* expression in wild-type seedlings. Relative expression was normalized to *ACT2* and data are the mean ± S.E. of n = 3 biological replicates.

**Supplementary Figure 2.**
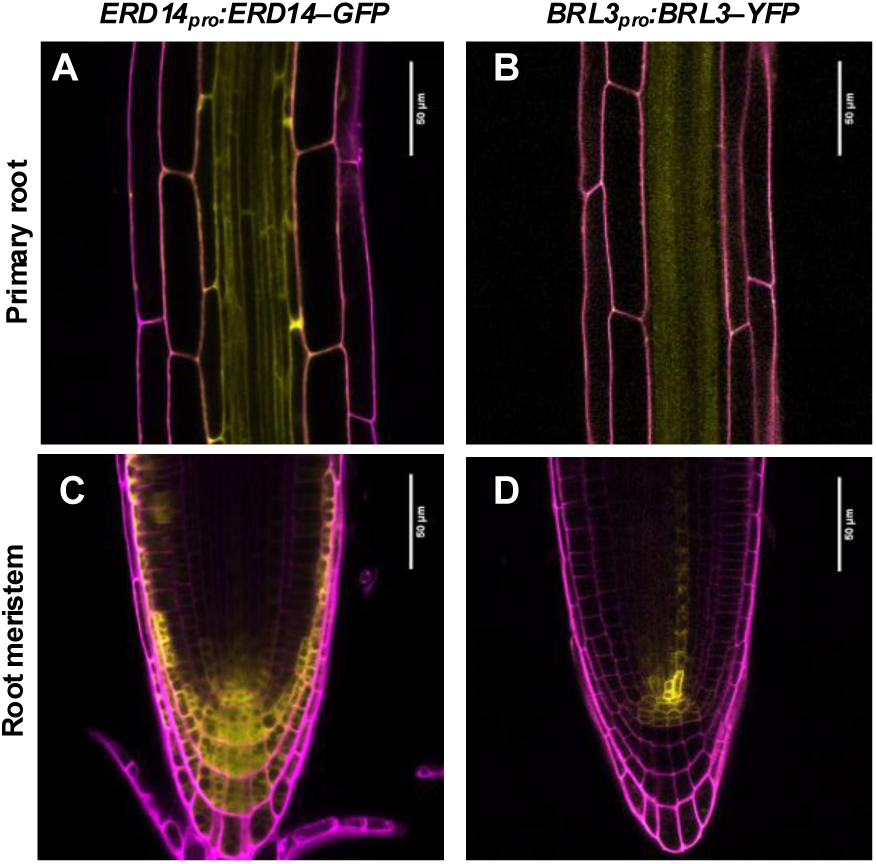
ERD14–GFP colocalizes with BRL3–YFP in different tissues. **A,B** Confocal microscopy of vascular-tissue detail in primary roots of transgenic lines expressing *ERD14_pro_:ERD14–GFP* and *BRL3_pro_:BRL3–YFP*. GFP/YFP are false-colored yellow and propidium-iodide counter staining is false-colored magenta. Scale bar: 50 µm. **C,D** Confocal microscopy of vascular-tissue detail in root meristems of transgenic lines expressing *ERD14_pro_:ERD14–GFP* and *BRL3_pro_:BRL3–YFP* constructs. GFP and YFP are false-colored yellow and propidium-iodide counter staining is false-colored magenta. Scale bar: 50 µm.

**Supplementary Figure 3.**
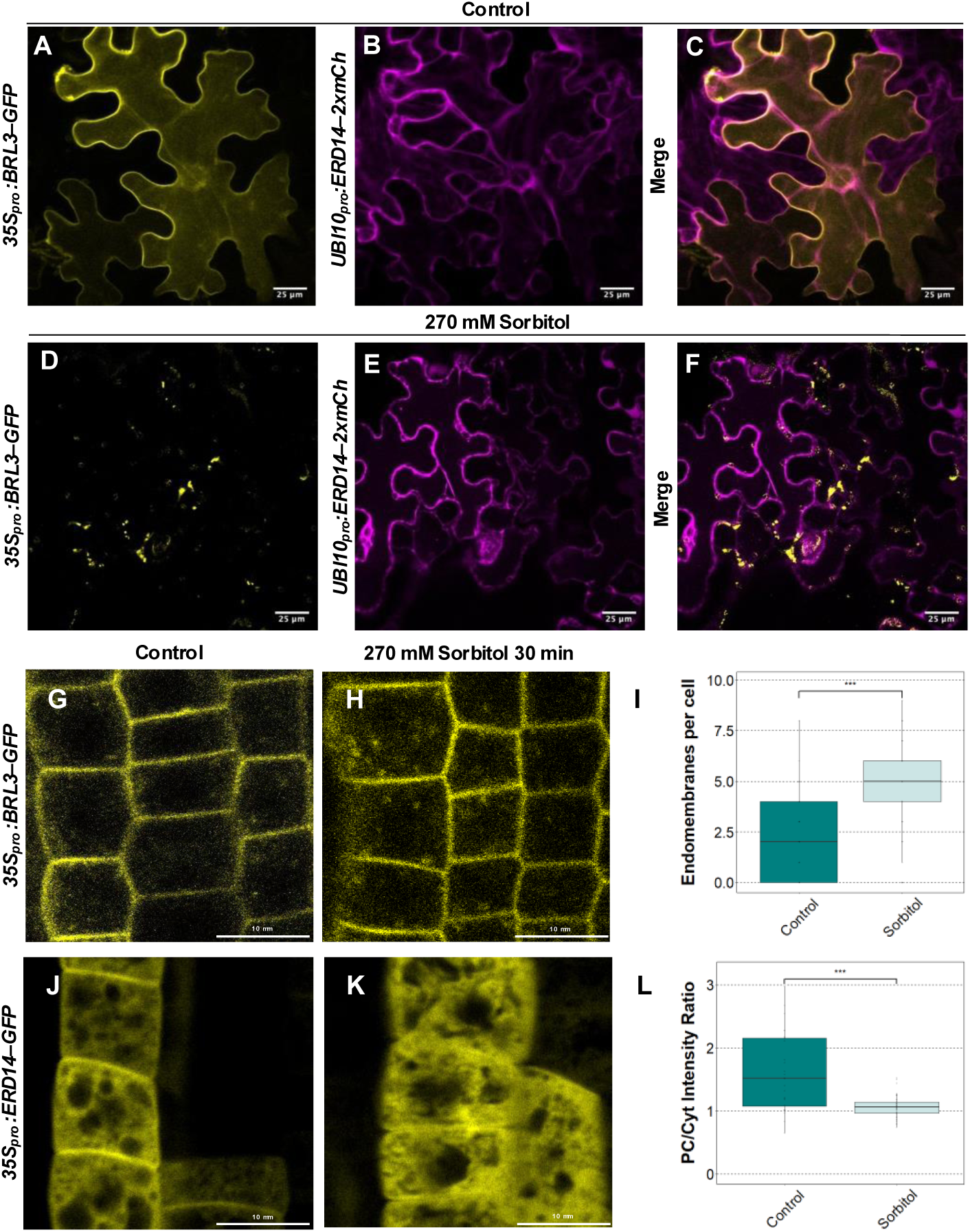
ERD14–mCherry colocalizes at the plasma membrane with BRL3–GFP and aggregates in endomembranes upon sorbitol treatment. **A–F** Confocal microscopy of *N. benthamiana* leaves transiently expressing *35S_pro_:BRL3– GFP* and *UBI10_pro_:ERD14–2xmCherry* constructs in control conditions (**A*–*C**) and upon 270 mM sorbitol treatment (**D–F**). GFP fluorescence is false-colored yellow and mCherry fluorescence in magenta. Scale bar: 25μm. **G–I** Confocal microscopy of Arabidopsis *35S_pro_:BRL3–GFP* root epidermis in control conditions (G) and after transfer to 270 mM sorbitol for 30 min (H). Scale bar: 10 µm. Quantification of endomembranes present per cell in *35S_pro_:BRL3–GFP* under control conditions and after 270 mM sorbitol for 30 min (I). Error bars depict the ± S.E. of the mean. Data are n = 86 and 125 cells for control and sorbitol respectively. Statistical significance is shown by asterisks following a Wilcoxon test (P < 0.05). **J–L** Confocal microscopy of Arabidopsis *35S_pro_:ERD14–GFP* root epidermis in control conditions (J) and after transfer to 270 mM sorbitol for 30 min (K). Scale bar: 10 µm. Quantification of the ratio of ERD14–GFP fluorescence intensity between plasma membrane and cytosolic regions under control conditions and after 270 mM sorbitol for 30 min (L). Error bars depict the ± S.E. of the mean. Data are n = 28 and 39 cells for control and sorbitol respectively. Statistical significance is shown by asterisks following a Student’s t-test (P < 0.05).

**Supplementary Figure 4.**
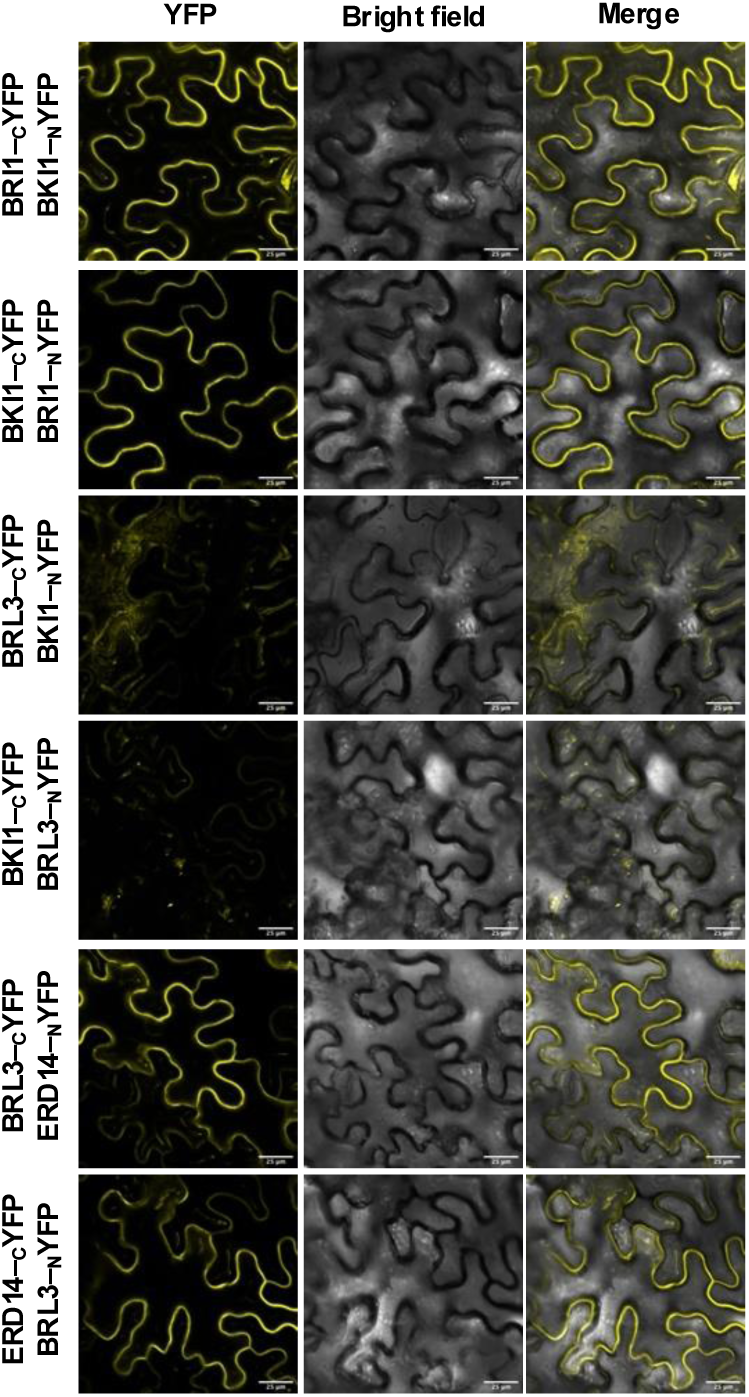
BiFC analysis of ERD14 interactions with BRL3. BiFC assay of BRL3*–*ERD14 physical interactions by transient expression in *N. benthamiana* leaf pavement cells. The known BRI1*–*BKI1 interaction was used as a positive control and BRL3*–*BKI1 non-interaction as a negative control. Scale bar: 25 µm.

**Supplementary Figure 5.**
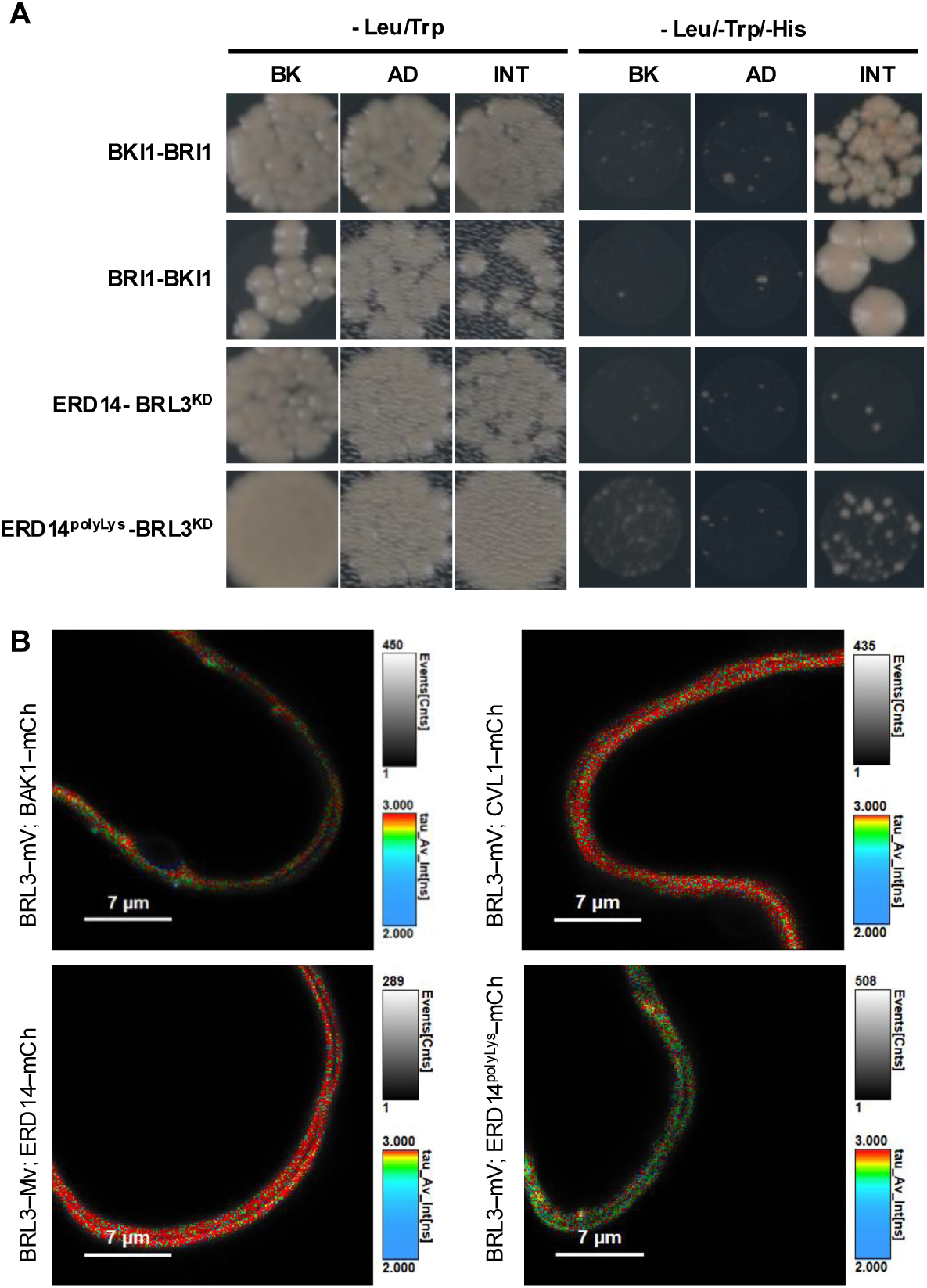
Analysis of the BRL3–ERD14 physical interaction by yeast two-hybrid and FRET–FLIM. **A** Yeast two-hybrid assay. Growth of clones harboring pGADT7-GW with an empty pGBKT7-GW (AD), pGBKT7-GW with an empty pGADT7-GW (BK), and the interaction between pGADT7-GW and pGBKT7-GW constructs (INT) on media without leucine and tryptophan (-Leu/-Trp, left) and media without leucine, tryptophan and histidine to test their interaction (-Leu/-Trp/-His, right). The known BRI1*–*BKI1 interaction was used as a positive control. **B** Raw FRET*–*FLIM images in which mCherry fluorescence is false-colored red and green for mVenus. Scale bar: 7 µm.

**Supplementary Figure 6.**
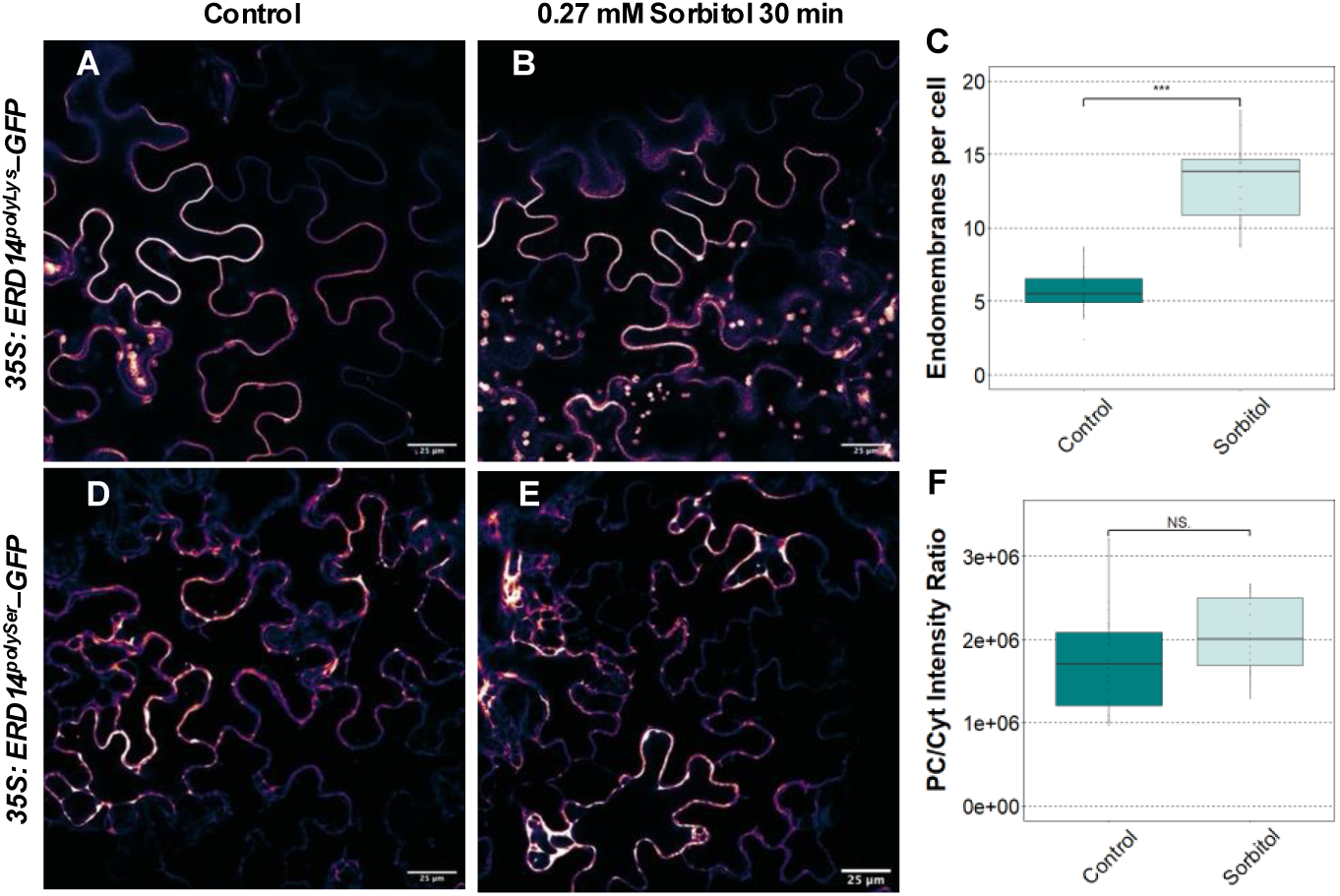
Analysis of ERD14^polyLys^ and ERD14^polySer^ dynamics upon osmotic stress. **A, B** Confocal microscopy of ERD14^polyLys^*–*RFP subcellular localization in transiently transgenic *N. benthamiana* leaves under control (A) and 270 mM sorbitol stress (B) conditions, respectively. Scale bar: 25 µm. **C** Quantification of ERD14^polyLys^*–*RFP endomembranes per cell as in A and B. Data are n > 15 cells. Error bars depict the ± S.E. of the mean. Statistical significance showed by asterisks after a Student’s t-test (P>0.05). **D, E** Confocal microscopy of ERD14^polySer^*–*RFP subcellular localization in transiently transgenic *N. benthamiana* leaves under control (D) and 270 mM sorbitol stress (E) conditions, respectively. Scale bar: 25 µm. **F** Quantification of ERD14^polySer^*–*RFP fluorescence intensity under control and 270 mM sorbitol stress as in D and E. Data are n > 15 cells. Error bars depict the ± S.E. of the mean. Statistical significance is shown by asterisks after a Student’s t-test (P < 0.05).

**Supplementary Figure 7.**
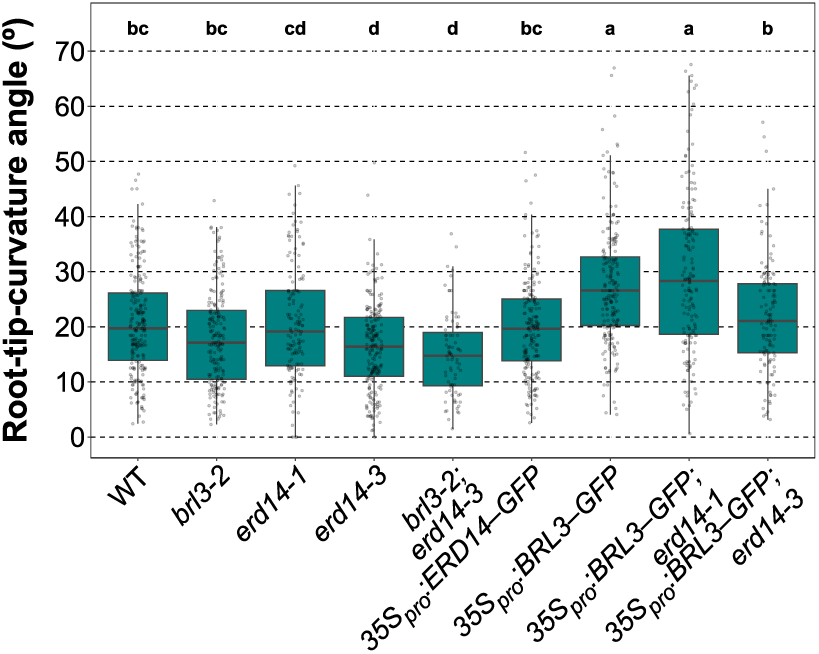
ERD14 is required for proper BRL3-mediated hydrotropic response. Root-tip angle after 24 h of sorbitol-induced osmotic stress. Data are n > 79 roots from 10 independent replicates. The effect of genotype on root-tip angle was determined by one-way Kruskal–Wallis test plus Bonferroni correction (P < 0.05).

**Supplementary Figure 8.**
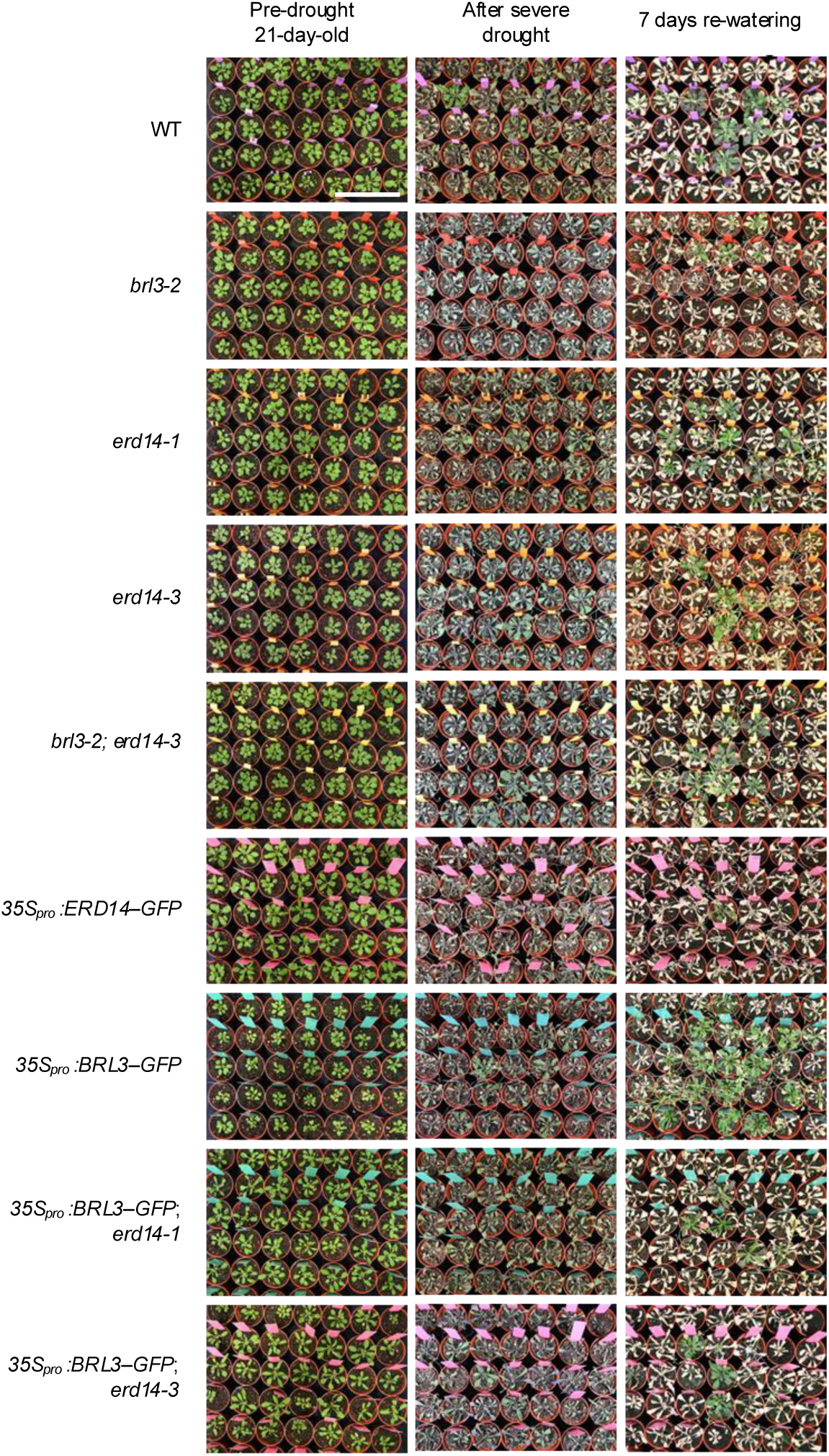
*35S_pro_:BRL3–GFP* resistance to severe water deprivation is reverted in *erd14* mutant backgrounds. Phenotypes of 40-d-old plants after 12 d of drought stress then 7 d of re-watering. Scale bar: 10 cm.

**Supplementary Figure 9.**
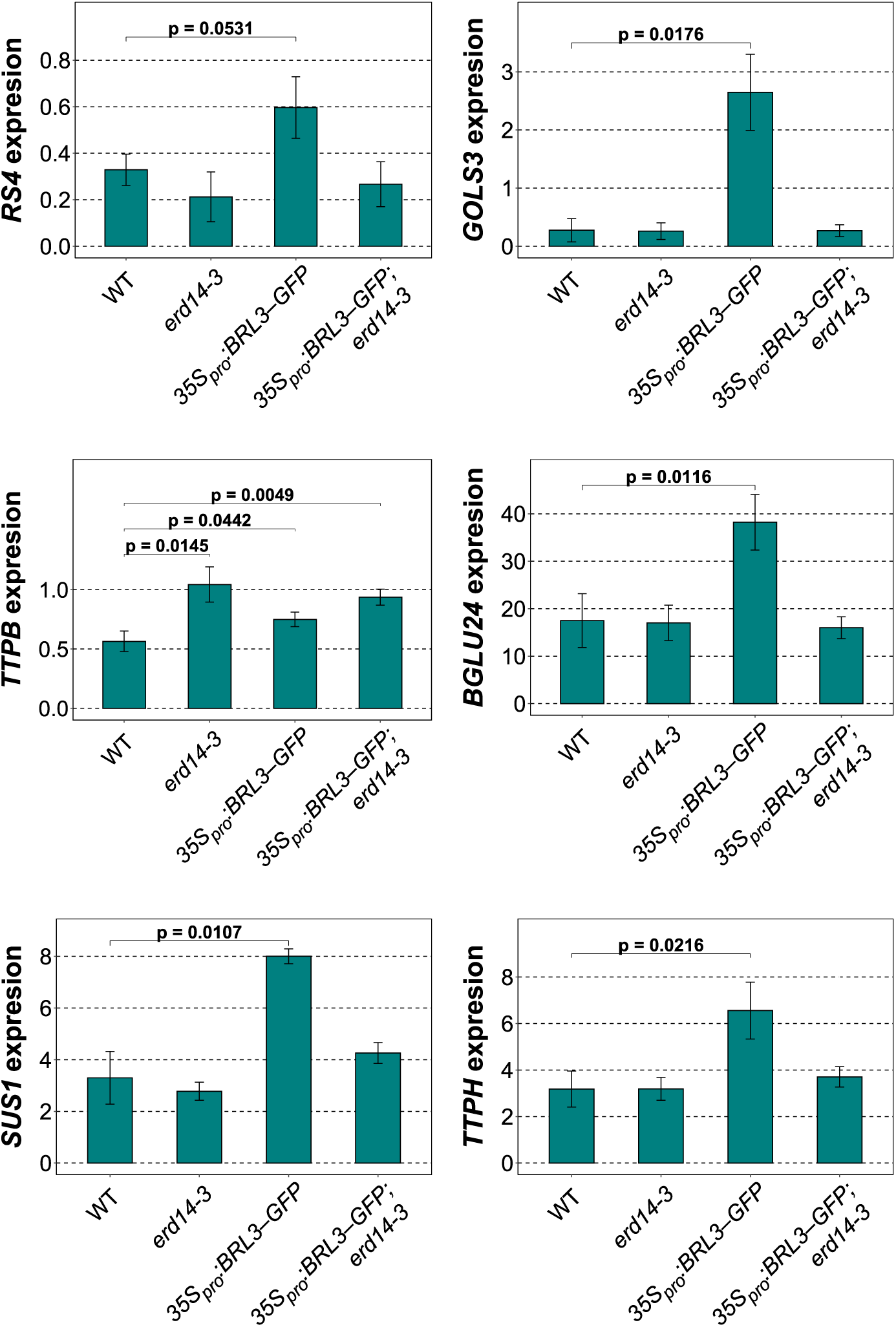
Select genes involved in osmoprotectant biosynthesis are differentially expressed in *erd14-3* and *35S_pro_:BRL3–GFP* backgrounds. RT–qPCR of transcripts encoding enzymes involved in biosynthesis of select osmoprotectant metabolites. Data are the mean ± S.E. of n = 3 biological replicas and relative expression was normalized to *ACT2*. The effect of genotype on means of expression levels was determined by Welch’s t-test.

---

**Supplementary Table 1.**
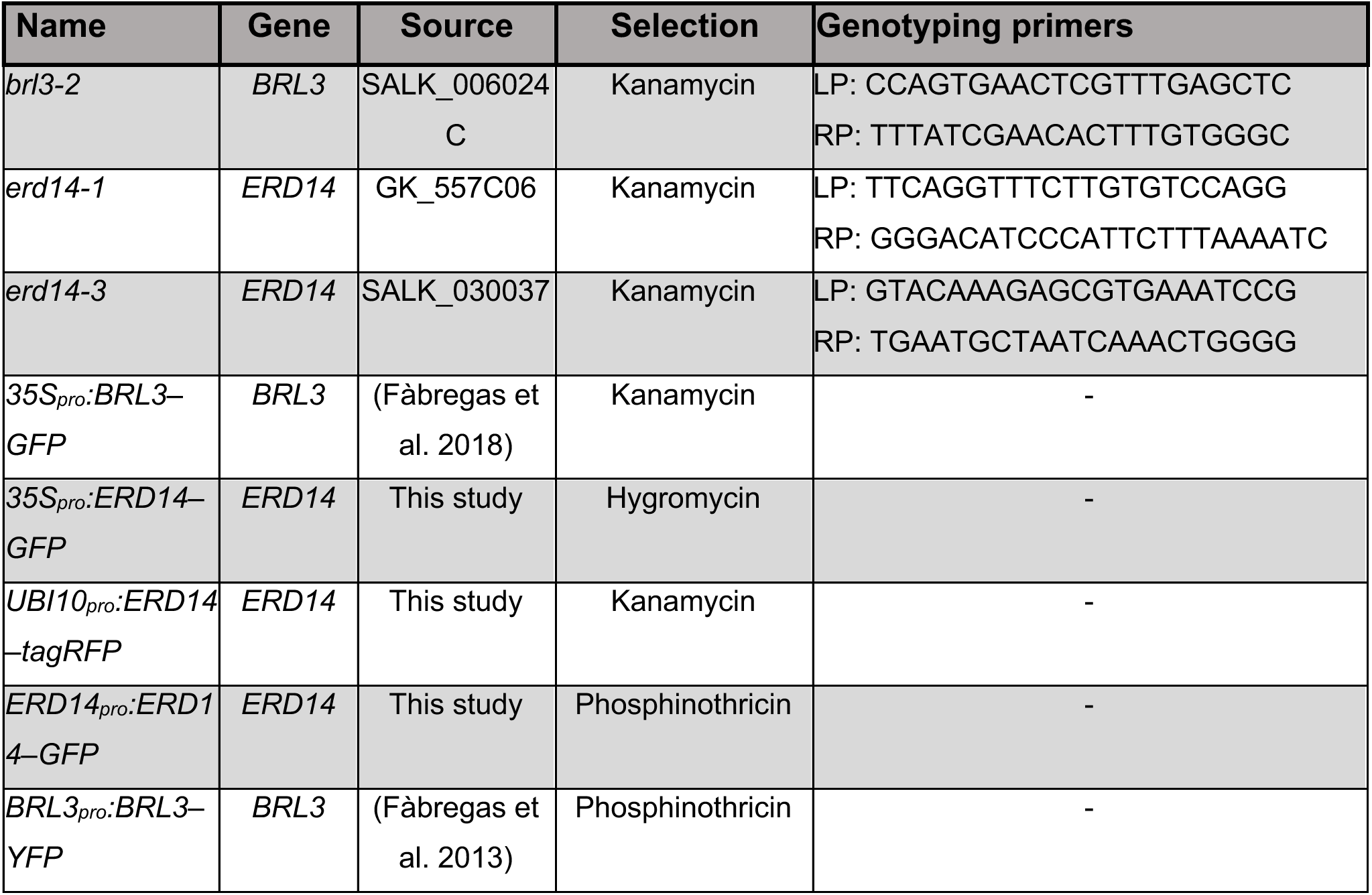
Plant material used and generated in this study.

**Supplementary Table 2.**
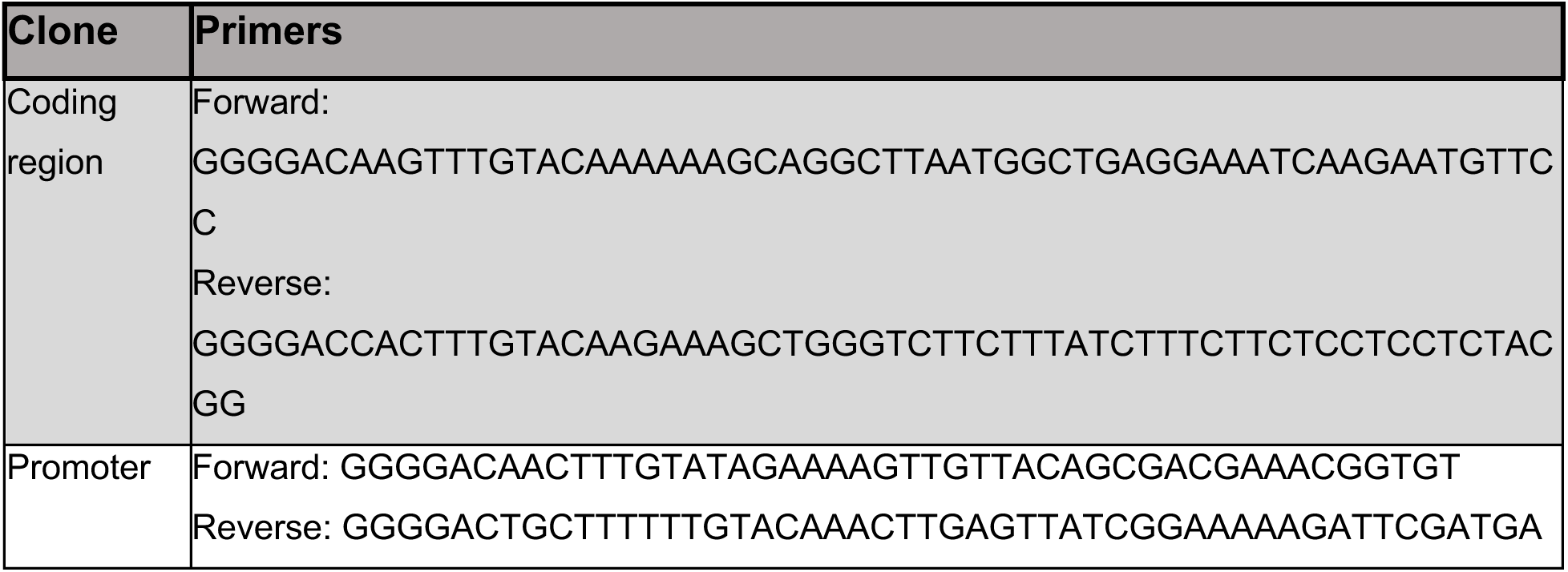
Primers used for cloning *ERD14* constructs.

**Supplementary Table 3.**
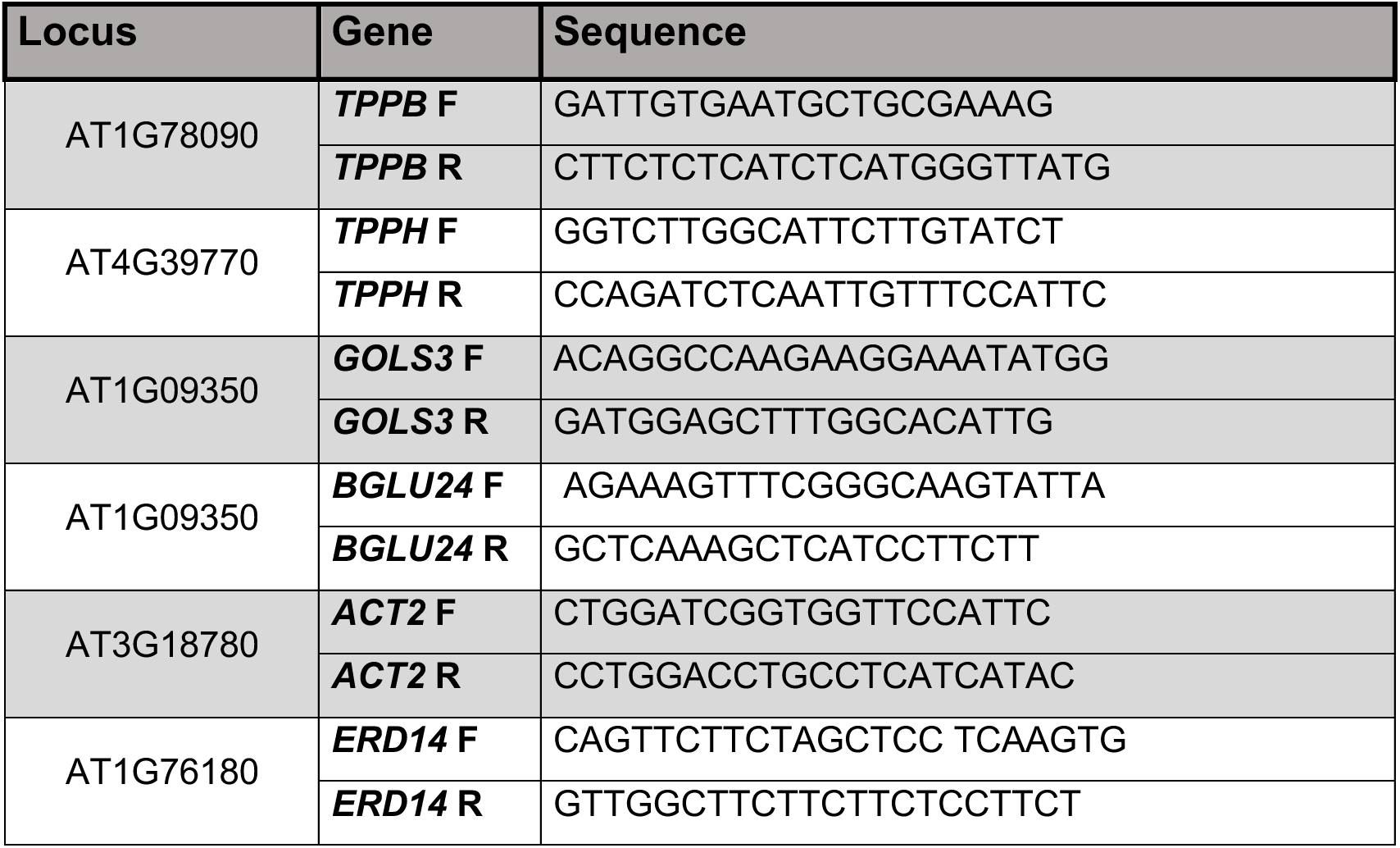
Primers used for RT–qPCR.

